# Capsular Polysaccharide Safeguards a Prophage–Bacterium Symbiosis by Preventing Collateral Attack and Promoting Viral Transmission

**DOI:** 10.64898/2026.09.09.750276

**Authors:** Mirjam Zünd, Lizett Ortiz de Ora, Nancy Haro-Ramirez, Elizabeth T Wiles, Jack T Leonard, Yuxin J Chen, Maria E Gallardo, Kenny Ho, Thomas P O’Malley, Shane Gonen, Matthew E Griffin, Katrine L Whiteson, Travis J Wiles

## Abstract

Many symbioses exist along a continuum from cooperation to conflict. Latent bacterial viruses known as prophages embody this duality. Acting as both partner and predator, they can enhance bacterial fitness while retaining the capacity to kill their hosts through lytic replication. Yet what determines the balance between cooperation and conflict in prophage–bacterium symbioses, and how these associations avoid collapse, remains poorly characterized. To identify mechanisms that stabilize prophage–bacterium partnerships, we used experimental evolution to perturb a natural association through repeated cycles of transmission and reinfection. This perturbation consistently selected mutant hosts lacking capsular polysaccharide production and exposed a hidden conflict that proved detrimental to both partners. During lytic outbreaks, host cells suffered lethal collateral prophage attack while dispersing virions became entrapped on neighboring cells and lysis debris, severely restricting transmission. Genetic and imaging-based studies revealed that capsular polysaccharides suppress these maladaptive interactions by limiting phage readsorption at the cell surface. We term this host-mediated safeguard the Hyperion effect, after the Greek Titan of light. Hyperion interactions enable phages to radiate outward from host populations, thereby averting collateral attack and promoting viral transmission. Our findings demonstrate that conflicts between prophages and their hosts can extend beyond individual cells to whole populations, with consequences that scale to shape patterns of prophage spread and microbial community assembly. More broadly, our work illustrates how mutually beneficial prophage–bacterium interactions can arise not only through cooperation, but also through the suppression of mutually detrimental conflicts.

## INTRODUCTION

Phages are often viewed as lethal predators of bacteria, yet this captures only part of their ecological role (1–3). Across microbiomes, phages shape bacterial communities not only through killing but also by driving gene flow and diversification, altering metabolism, and modulating nutrient cycling (1). These activities blur the boundary between antagonism and mutualism, highlighting how interactions that appear parasitic can, under certain conditions or across biological scales, support the persistence of both partners (2–5). However, the mechanisms that determine when phage–bacterium symbioses are defined by conflict or cooperation—and whether such relationships persist or collapse—remain poorly understood.

Temperate phages exemplify the tension between cooperation and conflict in phage– bacterium interactions. Following infection, temperate phage genomes can integrate into the bacterial chromosome, where they persist as vertically inherited prophages. In this lysogenic state, prophages often confer fitness-enhancing traits to their hosts, but they also represent latent internal threats. In response to host or environmental cues, including genotoxic stress, prophages can be induced to enter the lytic cycle, producing viral progeny that are released through host cell lysis (6). Consequently, the benefits of lysogeny come with the continual risk of lethal antagonism that can destabilize the association.

From an evolutionary perspective, the fitness costs imposed by prophages, together with selective pressures acting on bacterial populations, are expected to favor either the loss or inactivation of prophages over time. Consistent with this expectation, many bacterial chromosomes harbor degraded prophages, reflecting past associations that failed to persist (7, 8). Conversely, ecological conditions that favor phage spread may select for increased lytic replication and horizontal transmission, further destabilizing long-term associations (9, 10).

Whether driven by bacterial adaptation or phage transmission, both trajectories undermine the persistence of prophage–bacterium symbioses. Yet temperate phages remain widespread across bacterial lineages, with a substantial fraction of genomes encoding one or more intact prophages (6, 8). Their prevalence suggests that prophage–bacterium symbioses not only form readily but also endure despite selective pressures acting against them.

Explanations for the persistence of prophage–bacterium symbioses have largely focused on phage-encoded control over replication. For example, a recent quantitative model rooted in Parrondo’s paradox—in which alternating between two unfavorable strategies yields a net benefit—predicts that switching between lytic and lysogenic states stabilizes phage–host associations by limiting extreme population fluctuations (11). Moreover, comparative and experimental studies have revealed remarkable diversity in the replication control elements of temperate phages and show that these systems readily evolve in response to selection through changes in induction sensitivity and threshold (12–15). Together, these findings suggest a view of prophage–bacterium symbioses in which stability emerges through coevolutionary tuning of phage replication behaviors that balance lytic exploitation with long-term lysogenic persistence.

Here, we set out to link phage replication control directly to the stability of prophage– bacterium symbioses. Using an experimental evolution approach, we cycled a P2-like prophage through repeated rounds of induction and reinfection in its native *Escherichia coli* (*E. coli*) host, thereby driving the partnership beyond its co-adapted equilibrium. Rather than uncovering phage-encoded changes in replication control, this perturbation exposed a hidden source of self-destructive conflict in which endogenous prophages reattach to and kill their lysogen host population. This maladaptive state also impaired horizontal prophage transmission, as virions became entrapped on host cells and lysis debris. We found that host capsular polysaccharide expression prevents these mutually detrimental outcomes by masking a phage receptor on the bacterial cell surface. Finally, these interactions not only modulate the fate of individual prophage–host pairs at the cellular level, but also shape competitive fitness of populations within defined communities. Altogether, our findings suggest that bacterial cell-surface defenses against prophage progeny released from sister lysogens can determine the balance between conflict and cooperation, with consequences that scale from individual partnerships to microbial communities.

## Results

### Experimentally driving imbalance in a prophage–bacterium symbiosis

To probe mechanisms that balance prophage–host symbioses, we selected the P2-like temperate phage DuoHS and its native, human gut-derived *E. coli* HS host as a tractable model system. P2-like phages are tailed viruses belonging to the class Caudoviricetes and are widespread across lineages of Pseudomonadota (Proteobacteria) (15–17). DuoHS is stably integrated as a prophage at a defined chromosomal attachment site (*attB*) and becomes induced to lytically replicate in response to DNA damage and activation of the host SOS response. We previously characterized DuoHS replication and infection dynamics and established its amenability to genetic manipulation and single-virion imaging using fluorescence-based Phollow tagging (18). This model prophage–host system thus enables quantitative dissection of lytic and lysogenic replication and interactions across molecular, cellular, and population scales.

To drive imbalance in the DuoHS–HS symbiosis, we serially passaged DuoHS through alternating rounds of lytic and lysogenic replication on a susceptible *E. coli HS* host cured of the DuoHS prophage and carrying a restored *attB* integration site (Fig. 1A, B). To induce lytic replication, we used the genotoxic drug mitomycin C (MMC). Serial passaging is a classic strategy for enriching viral variants with enhanced infectivity, favoring traits that promote more efficient induction, host-cell attachment, establishment of infection, or increased virion production (19–23). We hypothesized that imposing this pressure would push the DuoHS–HS association beyond its apparently co-evolved relationship, shifting the balance toward increasingly antagonistic phage behaviors (9, 10) while exposing the mechanisms that normally constrain them.

**Figure 1.**
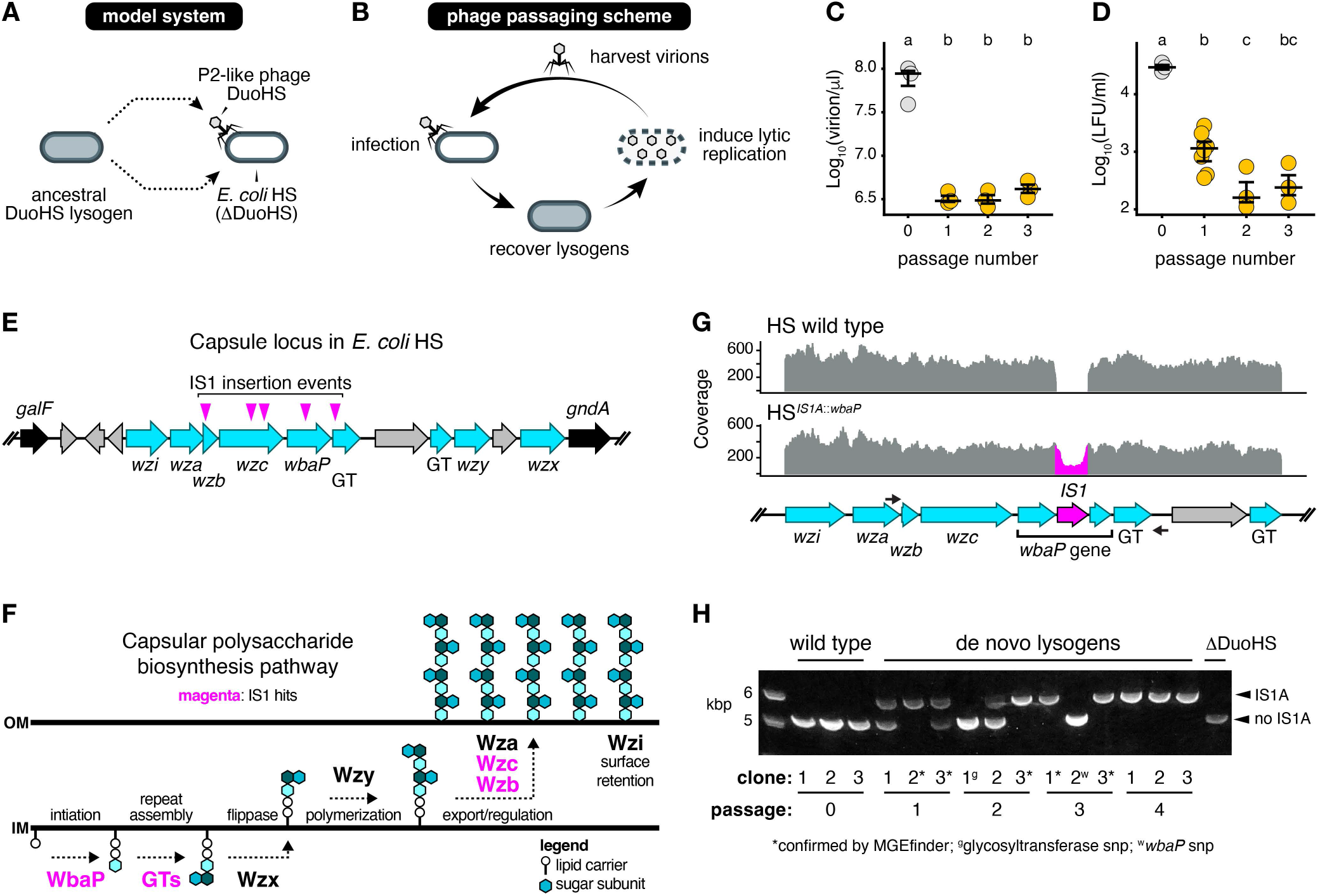
Serial passaging of DuoHS reduces virion abundance and lysogenic infectivity while selecting for host capsule mutations. **(A)** Model system: DuoHS was induced from an *Escherichia coli* HS lysogen and used to infect an engineered HS strain lacking the DuoHS prophage (ΔDuoHS). **(B)** Phage passaging scheme: DuoHS was serially passaged through repeated rounds of lytic and lysogenic replication. During each round, de novo lysogens were isolated, archived, and induced to produce virions for the subsequent passage. **(C)** Virion yield and **(D)** lysogen formation units (LFU) were quantified at each passage by qPCR and plating-based methods, respectively. Grey circles represent data derived from ancestral lysogens (passage 0), whereas amber circles represent data derived from lysogens archived at the indicated passages. **(E)** Organization of the capsular polysaccharide (CPS) biosynthesis locus in *E. coli* HS. Cyan arrows denote genes predicted to function in CPS production, while grey arrows denote unclassified or hypothetical genes. Magenta arrowheads indicate independent IS1 insertion events within DuoHS lysogens. **(F)** Predicted pathway for CPS biosynthesis in *E. coli* HS. CPS synthesis is initiated by WbaP- mediated transfer of the first sugar to a lipid carrier, followed by repeat-unit assembly by glycosyltransferases (GTs), translocation across the inner membrane (IM) by Wzx, polymerization by Wzy, and export by the Wza-Wzc complex prior to surface retention by Wzi. Genes disrupted by IS1A insertion are highlighted in magenta. **(G)** Read alignments confirming IS1A transposition into the *wbaP* gene of clone 2 from passage 1 (bottom) and its absence in the ancestral wild-type strain (top). IS1A insertion disrupts *wbaP*, creating a truncated open reading frame that terminates within the 5′ region of the IS1A element. The two black arrows indicate the binding site of the primer used to screen for IS1A mediated locus disruption shown in panel H. **(H)** PCR-based screen of the CPS locus in ancestral wild-type lysogens, ΔDuoHS, and de novo lysogen clones recovered from each passage. A ∼5 kb amplicon indicates the wild-type locus, whereas IS1A insertion produces a ∼5.78 kb amplicon. Double bands indicate mixed populations containing cells with and without IS1A insertions. Asterisks denote IS1 insertions confirmed by MGEfinder; ^g^ denotes a glycosyltransferase SNP and ^w^ denotes a *wbaP* SNP. Bars in each plot indicate the median and interquartile range. Statistical groupings denoted by different letters (p < 0.05) were determined by one-way ANOVA followed by Tukey’s HSD post hoc test for multiple comparisons.

### Serial passaging of DuoHS reduces extracellular virion abundance and lysogenic infectivity and selects for host mutations in capsular polysaccharide synthesis genes

Across serial lytic–lysogenic passages, we observed a rapid decline in measures of DuoHS replication and infectivity. The abundance of extracellular DuoHS virions following lytic induction, quantified by qPCR, decreased by more than an order of magnitude by passage two (Fig. 1C). Lysogen-forming units (LFUs), which indicate successful infection and establishment of lysogeny in new hosts, declined even more sharply, falling by nearly two orders of magnitude in response to passaging (Fig. 1D). To determine whether these unexpected outcomes of viral attenuation depended on features of the initial passaging scheme, we performed a second experiment in which we standardized multiplicity of infection, shortened induction and infection intervals, and removed antibiotics during lytic replication (Fig. S1A). Under these tightly controlled conditions, extracellular virion abundance and LFUs again declined rapidly (Fig. S1B,C), demonstrating that the phenotypic changes we had observed are a robust and reproducible consequence of serial passaging of DuoHS on its native host.

To identify the genetic basis underlying the apparent attenuation of DuoHS infectivity, we sequenced de novo lysogens isolated from each round of passaging. Surprisingly, we did not detect mutations within the integrated DuoHS prophage genome. Initial variant calling instead identified a single recurrent missense mutation in the host gene *pdeI*, which encodes a cyclic di-GMP phosphodiesterase (Fig. S2A). However, an in-frame deletion of *pdeI* in the ancestral *E. coli* HS background failed to recapitulate the observed defect in DuoHS lysogenic infectivity (Fig. S2B).

We therefore examined whether de novo lysogens harbored structural variants not readily detected by standard short-read alignment pipelines. Using MGEfinder, we found that the majority of isolates carried IS1A insertions within a bacterial chromosomal locus encoding a Wzx/Wzy-dependent capsular polysaccharide biosynthesis pathway (Fig. 1E) (24). IS1A insertions were distributed across four genes—encoding the initiating glycosyltransferase WbaP, a predicted glycosyltransferase, the phosphotyrosine phosphatase Wzb, and the tyrosine kinase Wzc—which together contribute to repeat unit assembly, polymerization, and surface export of capsular polysaccharides (Fig. 1F) (25–27). Disruption of any one of these genes is expected to abolish or substantially alter capsule production.

Long-read genome assembly of a representative de novo lysogen confirmed IS1A insertion within *wbaP* (HS*^IS1A^*^::*wbaP*^), an event predicted to disrupt WbaP function and likely perturb transcription of downstream capsule biosynthesis genes (Fig. 1G). PCR-based screening further revealed that 10 of 12 de novo lysogens from across all passages produced an expanded amplicon consistent with IS1A insertion (Fig. 1H). Multiple isolates yielded mixed PCR products, suggesting that these clones represent heterogeneous populations. Consistent with this observation, we identified low-frequency, independent single-nucleotide disruptions in the same capsular polysaccharide biosynthesis genes that are targeted by IS1A (Table S1). Notably, the only two de novo lysogens lacking detectable IS1A insertions instead harbored single base pair deletions in *wbaP* or the predicted glycosyltransferase gene (Fig. 1H, Fig. S2A). Similar patterns of IS1A mobilization and mutation were also recovered in the replicate passaging experiment (Table S1).

IS elements are abundant in *E. coli* and are well known to generate phenotypic diversity by disruption of surface polysaccharide loci (28, 29). We identified 24 IS1 elements in the *E. coli* HS genome, which ISfinder classified into three distinct isoforms, with IS1A being the most numerous (Fig. S2C and Table S2) (30). Given this extensive reservoir of endogenous IS elements, the repeated recovery of IS1A insertions in capsular polysaccharide biosynthesis genes during serial passaging indicates that DuoHS selects host subpopulations generated by IS1 mobilization and spontaneous mutations, rather than driving adaptive evolution of the phage itself.

### *E. coli* HS encodes a *Klebsiella*-like K47 capsule that restricts DuoHS infection

The clustering of IS1A insertions within capsular polysaccharide biosynthesis genes was unexpected, as *E. coli* HS was originally reported to encode an O9 O-antigen, without any prior annotation or phenotypic evidence of a capsular (K) antigen (31). Comparative genomic analysis of the HS capsule locus, however, revealed a high degree of homology to the K47 capsule of *Klebsiella* species, explaining why it was left unannotated using *E. coli*-focused databases (Fig. S2D). In line with this observation, it was recently proposed that the K47 capsule region and the adjacent O9 O-antigen locus were co-transferred from *Klebsiella* to *E. coli* HS in a single horizontal gene transfer event (32).

In *Klebsiella*, capsule expression can be associated with a pronounced surface polysaccharide layer (33) while K47 capsule has been linked to immune evasion and is frequently linked to virulent, carbapenem-resistant lineages (34, 35). In contrast, *E. coli* HS does not display an obvious capsular layer by transmission electron microscopy and lacks overt pathogenic or drug-resistant phenotypes (Fig. S2E,F) (36–38). This discrepancy suggests that HS may produce surface polysaccharides but lacks the extensive polymerization and surface retention required to generate a prominent extracellular capsule characteristic of prototypic encapsulated strains.

We reasoned that chemical staining might provide a more sensitive way of detecting surface-associated polysaccharides that are not readily visualized by electron microscopy. We therefore leveraged the ability of capsular polysaccharides to retain polyphenolic mordants and cationic dyes such as methylene blue (39). Wild-type HS cells displayed moderate to strong blue staining, consistent with the presence of a surface-associated polysaccharide layer, whereas HS*^IS1A^*^::*wbaP*^ cells stained pink, consistent with disruption of polysaccharide biosynthesis at its initiating step (Fig. 2A). To directly test whether this staining phenotype depends on the IS-targeted capsule genes, we deleted all four capsule genes identified in our passaging experiment (creating HS ΔCPS). Staining of this mutant strain recapitulated the polysaccharide-negative pink staining observed with HS*^IS1A^*^::*wbaP*^ (Fig. 2A). Conversely, inducible expression of the deleted capsule genes in the HS ΔCPS background restored the exopolysaccharide-positive blue staining characteristic of wild-type HS (Fig. 2A and Fig. S3A). Together, these results demonstrate that methylene blue staining phenotypes depend on genes encoding core components of the Wzx/Wzy-dependent capsular polysaccharide assembly pathway.

**Figure 2.**
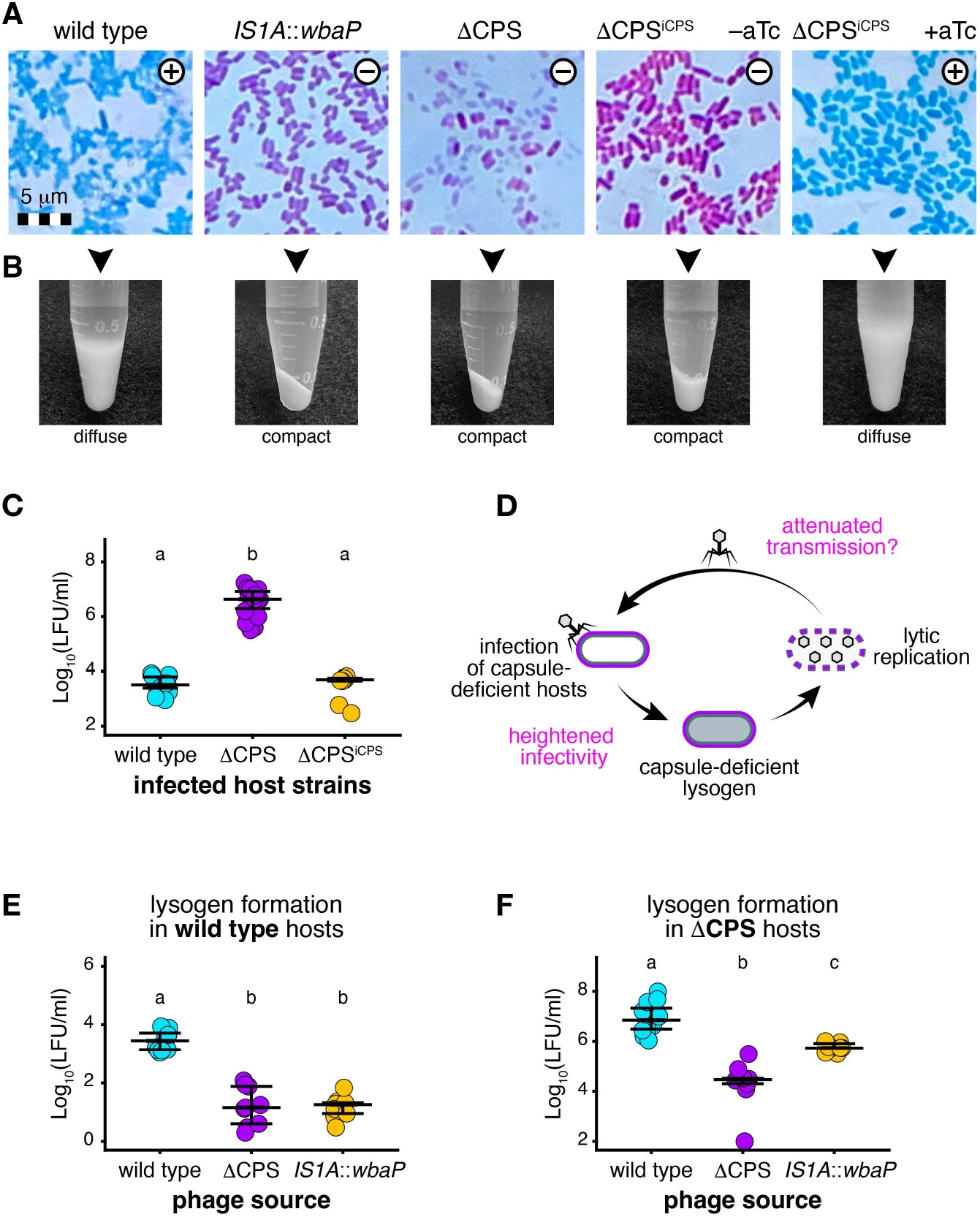
*E. coli* HS encodes a capsule that modulates DuoHS infection and transmission. **(A)** Differential staining phenotypes associated with capsule production. Wild-type HS, HS*^IS1A^*^::*wbaP*^, HS ΔCPS, and HS ΔCPS carry inducible CPS genes (iCPS) were stained with methylene blue. HS ΔCPS^iCPS^ was stained in the presence or absence of anhydrotetracycline (aTc) induction. Capsule- positive cells stain blue (⊕), whereas capsule-deficient cells stain pink (⊖). **(B)** Capsule-dependent pelleting phenotypes. Capsule-positive strains formed diffuse, poorly compacted pellets following centrifugation, whereas capsule-deficient strains formed compact pellets. **(C)** Lysogenic infectivity of DuoHS in target strains differing in capsule production. Lysogen formation units (LFU) were quantified following infection of wild-type (n = 15), ΔCPS (n = 23), and induced capsule-complemented ΔCPS^iCPS^ target strains (n = 9). **(D)** Proposed model for infection and transmission dynamics in capsule-deficient hosts. Loss of capsule production (purple cell outline) increases susceptibility to DuoHS infection and lysogenization but is possibly associated with reduced transmission of progeny phage following induction. **(E, F)** Lysogenic infectivity of DuoHS produced from different source lysogens. Phage induced from wild-type, ΔCPS, or HS*^IS1A^*^::*wbaP*^ lysogens (n = 9 each) was used to infect either wild-type target hosts (E) or ΔCPS target hosts (F). Lysogenic infectivity was quantified as lysogen formation units (LFU). Bars in each plot indicate the median and interquartile range. Statistical groupings denoted by different letters (p < 0.05) were determined separately for each plot using a Kruskal–Wallis test followed by pairwise Mann–Whitney U tests with Benjamini–Hochberg false-discovery-rate correction.

To further corroborate capsular polysaccharide production in HS using an independent physical readout, we exploited the highly hydrated and viscous nature of these surface- associated polymers, which can confer resistance to cell compaction during centrifugation. This property has been used as a qualitative indicator of capsule production in Wzx/Wzy-dependent systems in both *E. coli* and *Klebsiella pneumoniae* (40, 41). We found that wild-type HS cells formed diffuse, poorly compacted pellets following repeated low-speed centrifugation and wash cycles (Fig. 2B). In contrast, HS*^IS1A^*^::*wbaP*^ and HS ΔCPS strains formed tightly compacted pellets under identical conditions. Finally, forced expression of capsule genes in ΔCPS mutant increased resistance to compaction, yielding diffuse, wispy pellets (Fig. 2B). Together, these findings provide an independent physical correlate of surface polysaccharide production and further support the involvement of the identified Wzx/Wzy-dependent biosynthesis locus.

Having obtained evidence consistent with surface-associated capsular polysaccharide expression in *E. coli* HS, we next examined the functional consequences of capsule loss for DuoHS–HS interactions. The strong selection for non-encapsulated hosts during passaging indicated that such cells are hypersusceptible to DuoHS infection. Consistent with this idea, DuoHS lysogenic infection occurred at an approximately 1,000-fold higher frequency in HS ΔCPS cells than in wild-type hosts, whereas induced expression of the capsule genes reduced lysogenic infection and restored it to levels comparable to those observed in wild-type hosts (Fig. 2C). These results demonstrate that HS capsule expression restricts DuoHS infection. However, this effect was not evident from our original serial passaging experiments. Instead, the increased susceptibility of capsule-deficient hosts to DuoHS infection appeared to come at the expense of phage transmission (Fig. 2D). Supporting this inference, DuoHS derived either from HS*^IS1A^*^::*wbaP*^ or HS ΔCPS hosts had a marked reduction in lysogenic infectivity relative to phage induced from wild-type hosts, recapitulating the phenotype observed during passaging (Fig. 2E). Importantly, this defect persisted regardless of the recipient host genotype, indicating that the effect was determined by the source lysogen rather than the recipient host (Fig. 2F).

### Capsule governs DuoHS transmission and fitness through Kronos and Hyperion effects

One mechanism that could reconcile the opposing effects of capsule loss on DuoHS infectivity and transmission is altered virion adsorption. To investigate this possibility, we used fluorescently tagged Phollow phage virions to quantify phage attachment to wild-type and ΔCPS target cells (Fig. 3A). After 15 min of exposure, ΔCPS cells exhibited dramatically elevated phage binding, with nearly nine-fold more cells carrying at least one attached virion than wild-type cells (mean, 27.19% versus 3.07%; Fig. 3A,B). Moreover, an average of 8.1% of ΔCPS cells carried multiple attached virions, whereas multi-virion binding was not observed among wild-type hosts (Fig. 3A,B).

**Figure 3.**
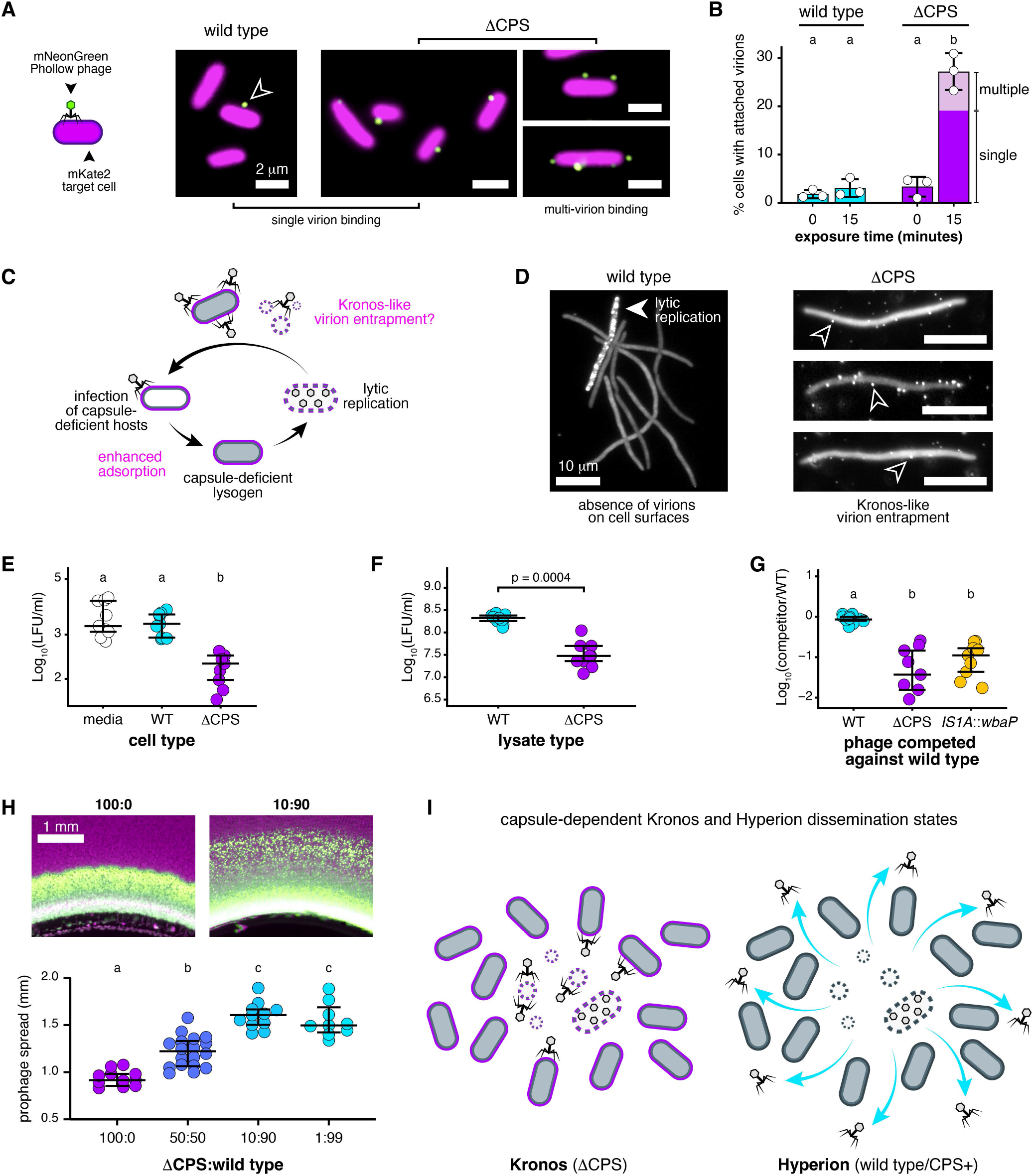
Lack of capsule leads to increased virion attachment and sequestration, limiting phage spread. **(A)** Representative images showing attachment of fluorescent DuoHS Phollow phage virions to wild-type (left) and ΔCPS (right) target cells. Target cells constitutively express mKate2-SpyCatcher (magenta), whereas DuoHS Phollow phage virions are labeled with mNeonGreen (green). The open arrowhead marks a virion attached to a wild-type cell. Representative examples of both single-virion and multi-virion binding events observed in ΔCPS populations are shown. **(B)** Percentage of wild-type and ΔCPS cells with at least one attached DuoHS virion at 0 and 15 min post-exposure. For ΔCPS populations at 15 min, the proportions of cells with single attached virions (dark purple) or multiple attached virions (light purple) are shown separately. Bars indicate means ± SD from three biological replicates (individual points), each containing 3–8 non-overlapping fields of view and 178–767 cells per replicate. **(C)** Proposed model in which enhanced DuoHS adsorption in capsule-deficient hosts promotes Kronos-like virion entrapment following lytic replication. **(D)** Representative images of DuoHS outbreaks in wild-type and ΔCPS Phollow lysogen populations. The white arrow indicates a lysogen undergoing induced lytic replication and virion production. Open arrowheads indicate virions attached to neighboring ΔCPS sister lysogens. **(E)** Infectious phage remaining in supernatants following pre-incubation with media alone (n = 9), wild-type cells (n = 9), or ΔCPS cells (n = 9), quantified as lysogen formation units (LFU). **(F)** Infectious phage quantified as LFU in the presence of cell-free lysate generated from wild-type (n = 9) or ΔCPS (n = 9) cells. **(G)** Head-to-head infection competition of phage populations derived from ΔCPS (n = 9) or HS*^IS1A^*^::*wbaP*^ (n = 9) lysogens against wild-type-derived phage (n = 9). Values represent the log^10^ transformed ratio of LFUs generated by each competitor relative to wild-type-derived phage following co-infection. **(H)** Representative images (top) showing dissemination of DuoHS virions carrying an mNeonGreen lysogeny reporter from a central agar plug into surrounding communities of mKate2-labeled target cells (magenta) containing the indicated ΔCPS:wild-type cell ratios. Newly lysogenized cells acquire green fluorescence, producing a visible prophage dissemination front. Plot (bottom) shows linear dissemination distances for each condition. 100:0 (n = 11), 50:50 (n = 18), 10:90 (n = 11), and 1:99 (n = 9). Individual points represent dissemination assays from six independent biological replicates per condition. **(I)** Model of capsule-dependent Kronos and Hyperion dissemination states. In the Kronos state (left), capsule-deficient host populations (purple outline) locally entrap DuoHS virions on neighboring cells and lysis debris, restricting dissemination. In the Hyperion state (right), capsule-expressing host populations (dark gray outline) preserve virion infectiousness and create corridors for phage dissemination, promoting long-range spread of infectious progeny (cyan arrows). Bars for panels E-H indicate the median and interquartile range. Statistical groupings denoted by different letters (p < 0.05) were determined separately for each plot. Panels E and G were analyzed using a Kruskal-Wallis test followed by pairwise Mann-Whitney U tests with Benjamini-Hochberg false-discovery-rate correction. Panels B and H were analyzed using one-way ANOVA followed by Tukey’s multiple-comparison test. Different letters indicate significant differences (p < 0.05). Panel F was analyzed using two-sided Mann-Whitney U tests.

The increased frequency of phage attachment to ΔCPS cells suggests that capsule expression limits access to a host determinant required for DuoHS adsorption, consistent with prior work indicating that P2-like phages recognize lipopolysaccharide (LPS)-associated surface structures (42). At the same time, the frequent accumulation of multiple virions on ΔCPS cells also suggests that increased adsorption may come at the expense of phage transmission.

Specifically, the heightened adsorption of DuoHS in ΔCPS populations may create a form of self-limiting phage loss analogous to the recently described *Kronos effect*—named after the Greek Titan who consumed his own offspring—whereby phage progeny become sequestered through reattachment to already infected host cells (Fig. 3C) (43). To probe whether capsule loss gives rise to virion fates consistent with Kronos-like interactions, we induced lytic outbreaks of in wild-type and ΔCPS Phollow lysogen populations and monitored the localization of newly released particles by live imaging. In ΔCPS populations, newly released phage progeny frequently landed on intact sister host cells, where multiple virions accumulated along cell surfaces (Fig. 3D). In contrast, surface-associated virions were rarely observed in wild-type populations, in line with our prior time-lapse imaging results (18).

To determine whether Kronos-like interactions measurably reduce the pool of infectious virions, we first tested whether capsule-deficient cells could function as phage sponges. We performed a pull-down assay by incubating DuoHS virions with wild-type cells, ΔCPS cells, or media alone before quantifying unbound infectious particles in the supernatant. Consistent with virion sequestration, pre-exposure to ΔCPS cells reduced LFUs by approximately one order of magnitude relative to both wild-type and media-only controls (Fig. 3E). We next asked whether cell lysis debris alone could have a similar effect. To test this, cell-free lysates generated from wild-type or ΔCPS hosts by inducible expression of phage λ lysis genes (see Methods and Fig. S3B–D) were incubated with DuoHS virions prior to LFU quantification. As with intact cells, exposure to ΔCPS-derived lysate significantly reduced LFUs relative to wild-type lysate (Fig. 3F), indicating that non-encapsulated host material released during lysis can sequester virions and/or disrupt productive receptor engagement. Finally, we tested whether Kronos-like interactions translated into reduced phage fitness, measured here as competitive transmission between hosts. We performed head-to-head competition assays using equal numbers of phage virions derived from wild-type, ΔCPS, or HS*^IS1A^*^::*wbaP*^ hosts. Phage populations originating from ΔCPS and HS*^IS1A^*^::*wbaP*^ hosts, which are produced in the presence of capsule-deficient lysis material, were strongly outcompeted by those derived from wild-type hosts (Fig. 3G).

Together, these experiments indicate that capsule loss promotes Kronos-like interactions that diminish the pool of infectious virions through local entrapment. This naturally raised the question of whether such interactions could also limit the net spread of transmitting virions through bacterial communities. To test this, we developed a virion dissemination assay in which the spread of infectious DuoHS particles could be directly visualized (see Methods). Briefly, DuoHS virions carrying an mNeonGreen lysogeny reporter were embedded within a central agar plug and allowed to disperse into a surrounding community of mKate2-tagged target cells. As virions spread from the point of inoculation, newly lysogenized cells acquire green fluorescence, generating a visible dissemination front that reports the net spatial spread of infectious phage particles (Fig. 3H, top). We hypothesized that communities composed entirely of ΔCPS cells would entrap virions and restrict dissemination, whereas increasing proportions of wild-type cells would create corridors for phage spread. In line with this prediction, host community composition strongly influenced the spatial dissemination of DuoHS progeny. Communities comprising ΔCPS cells alone limited viral spread from the inoculation site, whereas increasing proportions of wild-type cells progressively expanded dissemination fronts (Fig. 3H).

Altogether, these experiments reveal two distinct capsule-dependent dissemination states. Capsule-deficient HS populations are dominated by Kronos-like interactions that entrap DuoHS progeny, reducing infectiousness, fitness, and spatial dissemination. In contrast, capsule-expressing populations promote viral transmission and competitive fitness. We refer to this host-mediated dissemination state as the *Hyperion effect*, after the Greek Titan of light. Whereas Kronos interactions consume phage progeny through local entrapment, Hyperion interactions enable virions to radiate outward through bacterial populations and communities, promoting transmission and expanding viral range (Fig. 3I)

### Capsule shields host cell populations from lethal collateral prophage attack

Thus far, our results suggest that the Hyperion effect, mediated here through host capsule expression, provides a substantial benefit to DuoHS. At first glance, this benefit appears one-sided, favoring the prophage despite the considerable host investment in capsular polysaccharide production. However, because capsule is ultimately a host-encoded trait, we reasoned that it must also provide a selective advantage to the bacterium. We hypothesized that capsule expression acts to protect host populations from lethal outcomes associated with DuoHS outbreaks.

To probe this possibility, we used time-lapse microscopy to visualize induction of DuoHS in wild-type mNeonGreen Phollow lysogens and subsequent transmission into ΔCPS target cells (Fig. 4A). In this experimental setup, MMC-induced viral progeny released from wild-type hosts can be directly observed infecting nearby ΔCPS cells, where secondary lytic replication produces DuoHS Phollow phage virions tagged with mKate2. Previously, we showed that DuoHS can lytically replicate and spread among wild-type, capsule-expressing *E. coli* HS cells, but at low frequency (18). In striking contrast, MMC-induced outbreaks produced extensive virion adsorption and widespread lytic replication in ΔCPS target populations (Fig. 4B, Movie S1). We also observed instances in which mass virion attachment was followed by host cell lysis without detectable phage production (Fig. 4C, Movie S2). These events are consistent with the phenomenon of “lysis from without”, in which high-multiplicity phage adsorption triggers cell lysis in the absence of productive phage replication (44, 45).

**Figure 4.**
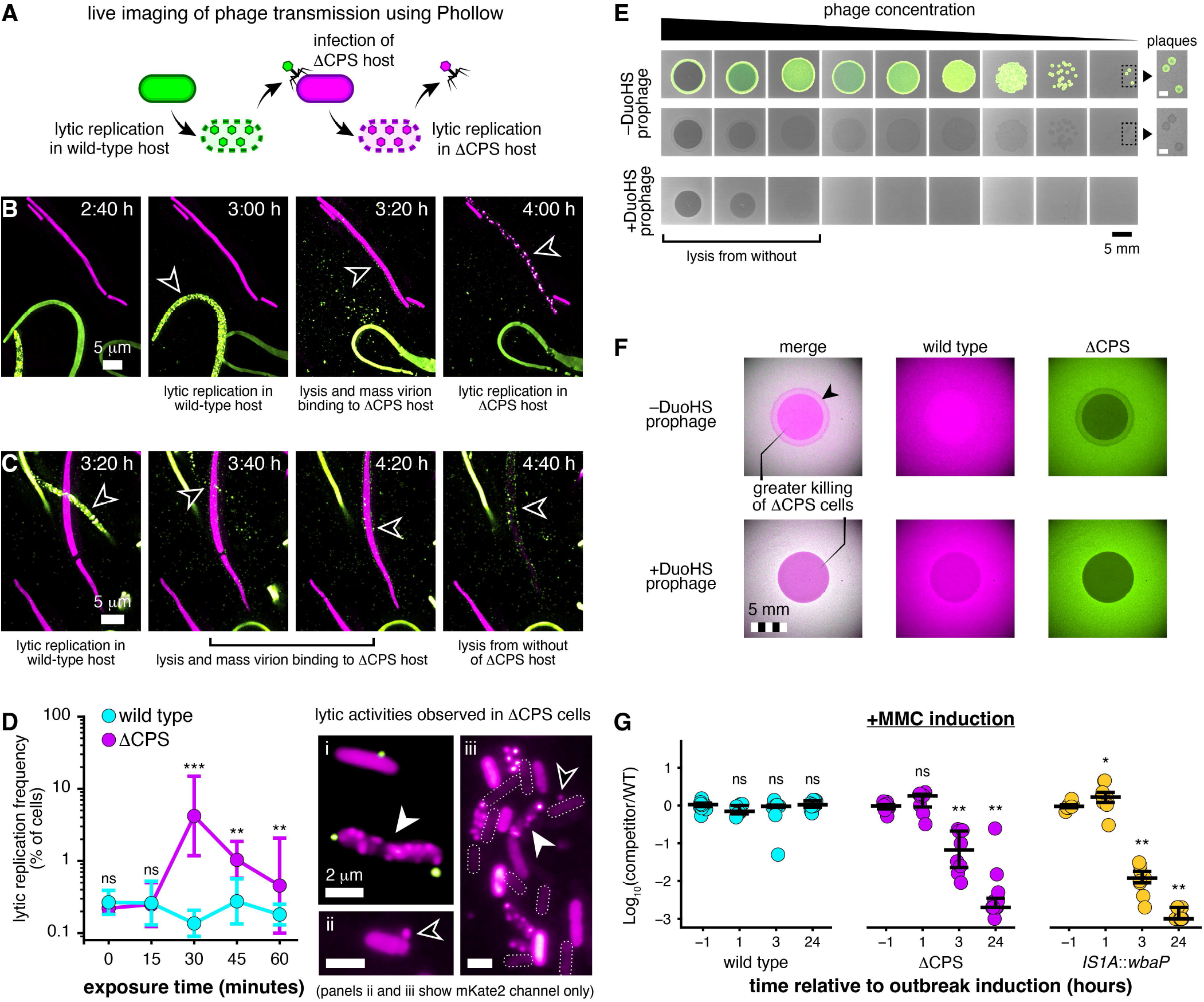
Capsule protects host cell populations from lethal phage interactions. **(A)** Schematic illustrating visualization of phage transmission using Phollow. Wild-type Phollow lysogens express an mNeonGreen (mNG)-SpyCatcher protein that labels DuoHS virions during lytic replication, producing green fluorescent Phollow phage virions. Following lysis, released virions infect and lytically replicate within ΔCPS target cells expressing mKate2-SpyCatcher, generating mKate2 Phollow phage progeny. **(B)** Time-lapse images showing productive DuoHS transmission from a wild-type Phollow lysogen to a ΔCPS target cell. Virion assembly is first observed within an mNG-labeled wild-type lysogen (3:00 h, arrowhead), followed by host lysis and mass virion adsorption to a neighboring mKate2-labeled ΔCPS target cell (3:20 h, arrowhead). Secondary lytic replication within the target cell produces mKate2 Phollow phage progeny (4:00 h, arrowhead). **(C)** Time-lapse images showing DuoHS-mediated lysis from without of a ΔCPS target cell. Following induced lytic replication in a neighboring wild-type lysogen, numerous released virions accumulate on the surface of a ΔCPS cell (3:20-4:20 h, arrowheads). This is followed by loss of cellular fluorescence and cell lysis (4:40 h, arrowhead) without detectable secondary phage production. **(D)** Lytic replication following exposure of wild-type or ΔCPS target populations to purified DuoHS Phollow phage virions. Left: Frequency of lytic replication over time. Points represent the geometric mean ± SD from the same imaging dataset shown in Fig. 3B, comprising three biological replicates, each containing 3–8 non-overlapping fields of view containing 178-767 cells. The limit of detection for lytic replication ranges from 0.13% to 0.56%. Data were analyzed using a binomial generalized linear mixed-effects model. Post hoc genotype comparisons at each time point were performed using estimated marginal means with FDR correction (*** < 0.001, ** = 0.004, ns = not significant). Right, representative images from ΔCPS populations. (i) Instance of productive lytic infection, indicated by assembly of mKate2 Phollow phage progeny within a cell carrying two attached mNeonGreen Phollow phage virions (white arrowhead). (ii) Reattachment of a newly released mKate2 Phollow phage virion (open arrowhead). (iii) ΔCPS cells 30 min post phage exposure undergoing lytic replication (white arrowhead), binding phage progeny (open arrowhead), or displaying reduced cytosolic fluorescence (dashed outlines), consistent with lysis from without-like events. For clarity, panels ii and iii show the mKate2 channel only. **(E)** Spot assays using DuoHS virions carrying an mNeonGreen lysogeny reporter. Phage spots represent 10-fold serial dilutions beginning with undiluted lysate. For ΔCPS target cell lacking a DuoHS prophage, the upper row shows fluorescence merged with brightfield, and the lower row shows brightfield alone. Newly lysogenized cells express green fluorescence. The bottom row shows an identical assay performed on ΔCPS lysogen carrying a DuoHS prophage. Insets show zoomed views of plaques from the boxed regions. White scale bars, 1 mm. **(F)** Representative mixed-lawn DuoHS DisCo assays containing mKate2-labeled wild-type (magenta) and mNeonGreen-labeled ΔCPS (green) populations either lacking (top) or carrying (bottom) a DuoHS prophage. Merged and individual fluorescence channels reveal depletion of ΔCPS cells relative to wild-type cells within the zone of phage exposure. Black arrowhead denotes a ring- like region enriched in ΔCPS cells that corresponds to elevated lysogeny observed in the spot assays shown in panel E. **(G)** Fitness consequences of capsule loss during DuoHS outbreaks. Relative abundance of ΔCPS and HS*^IS1A^*^::*wbaP*^ lysogens compared with wild-type lysogens before induction (−1 h) and throughout a phage outbreak. Panels show competitions between wild-type and wild-type lysogens (cyan, n = 9), ΔCPS and wild-type lysogens (purple, n = 9), and HS*^IS1A^*^::*wbaP*^ and wild-type lysogens (amber, n = 9). Relative abundance is expressed as Log^10^(CFU competitor/wild type). Bars indicate the median and interquartile range. Within each competition, time points 1, 3, and 24 were compared to time point -1 using a two-sided paired Wilcoxon signed-rank test with FDR correction ( * < 0.05, ** < 0.01, ns = not significant).

To determine whether the enhanced lytic activities observed in ΔCPS populations arose directly from phage transmission rather than the combined effects of MMC-induced stress, we returned to the purified virion infection assay used in Fig. 3. Here, Phollow was used to track DuoHS transmission as in Fig. 4A, but in the absence of MMC and co-cultured lysogens. Little to no lytic replication was observed in either wild-type or ΔCPS populations at 0 or 15 min following addition of DuoHS virions. However, by 30 min, a wave of lytic replication emerged in ΔCPS populations that persisted throughout the assay period, whereas lytic events remained at or below the limit of detection in wild-type hosts (Fig. 4D, left). mNeonGreen Phollow phage virions could be found attached to ΔCPS cells that simultaneously contained assembling mKate2 Phollow phage progeny (Fig. 4Di, white arrowhead). We also occasionally observed newly released mKate2 Phollow phage virions reattaching to ΔCPS cells, consistent with Kronos-like interactions (Fig. 4Dii,iii, open arrowheads). At the apparent peak of lytic replication (30 min post-exposure), many ΔCPS cells displayed reduced cytosolic fluorescence (Fig. 4Diii, cells with dashed outlines), reminiscent of the lysis from without events captured by time-lapse imaging in Fig. 4C. At the same time point, wild-type cells maintained robust fluorescent labeling (Fig. S4A). Together, these findings demonstrate that the heightened adsorption associated with capsule loss is sufficient to drive secondary lytic replication through phage transmission alone, independent of MMC-induced stress.

Building on these single-cell imaging observations, we next sought to characterize the balance between lethal and non-lethal outcomes associated with DuoHS outbreaks in ΔCPS populations. To do so, we performed spot assays using serial dilutions of DuoHS virions carrying an mNeonGreen lysogeny reporter (Fig. 4E, top). In these assays, green fluorescence marks non-lethal lysogenic infection, whereas zones of clearing indicate lethal outcomes arising from either lytic replication or lysis from without. Spotting DuoHS virions onto lawns of ΔCPS target cells lacking a DuoHS prophage produced extensive clearing accompanied by robust lysogeny reporter activity across a broad range of phage inputs. At lower phage concentrations, discrete plaques were readily observed, confirming that ΔCPS populations support productive lytic replication. In contrast, minimal clearing, and little to no lysogeny were observed in wild- type lawns even at the highest phage concentrations, consistent with capsule-mediated defense against DuoHS infection (Fig. S4B). To disentangle the relative contributions of lytic replication and lysis from without to lethal outcomes, we repeated the spot assay using ΔCPS cells carrying a DuoHS prophage. Because prophage-mediated immunity often protects lysogens from productive lytic reinfection (46), host killing within lysogen populations is expected to occur primarily through lysis from without at high multiplicities of infection. Consistent with this prediction, plaque formation was completely abolished while pronounced zones of clearing persisted at the highest phage concentrations (Fig. 4E, bottom). These results indicate that lethal interactions in ΔCPS populations can arise through either productive lytic replication or lysis from without, with the dominant mechanism determined by local phage density and prophage carriage.

Having established that capsule expression protects hosts from lethal DuoHS outcomes, we next asked whether it was sufficient to drive selection during prophage outbreaks within a community. To address this question, we employed a co-culture plaque assay inspired by the Phage DisCo method (47), in which DuoHS virions were spotted onto soft agar containing differentially labeled wild-type (magenta) and ΔCPS (green) populations. In mixed communities lacking a DuoHS prophage, wild-type cells persisted throughout the zone of phage exposure, whereas ΔCPS cells were markedly depleted (Fig. 4F, top). Notably, ΔCPS cells formed a distinct ring surrounding the central zone of clearance (Fig. 4F, top, black arrowhead). A similar pattern was observed in spot assays (Fig. 4E), where this region corresponded to lysogeny reporter activity, suggesting that local phage density might influence the balance between lytic and lysogenic outcomes. Importantly, the competitive advantage associated with capsule expression also manifested in lysogenic populations (Fig. 4F, bottom). Despite protection from productive lytic superinfection, ΔCPS lysogens were depleted to a much greater extent than wild-type lysogens within the zone of phage exposure. Capsule expression therefore provides a selective advantage during phage attack, favoring capsule-expressing hosts regardless of prophage carriage.

These observations led us to predict that capsule expression should be strongly selected for during active prophage outbreaks. To test this prediction, we competed wild-type lysogens against either a wild-type control, a ΔCPS lysogen, or an *IS1A*::*wbaP* lysogen and induced synchronized outbreaks with MMC. Prior to induction, all competitors were present at comparable frequencies (Fig. 4G, “-1”). Following MMC treatment, wild-type populations remained stable, whereas both capsule-deficient competitors declined sharply, falling by several orders of magnitude within 24 h (Fig. 4G). In contrast, no competitive differences were observed in the absence of MMC induction (Fig. S4C), demonstrating that fitness defects were not due to intrinsic growth differences. Capsule expression is therefore strongly selected for during DuoHS outbreaks, protecting lysogenized hosts from collateral attack by prophage progeny released from sister cells.

### Capsule restricts DuoHS infection by masking an O9 O-antigen receptor

The pronounced sensitivity of capsule-deficient hosts to DuoHS enabled the identification of the bacterial receptor required for infection. As noted above, receptors for P2-like phages are generally associated with LPS, but the precise structure engaged by DuoHS has not been defined. We therefore used a conventional enrichment approach to isolate host variants resistant to DuoHS-mediated killing. Exposure of ΔCPS populations to high densities of DuoHS virions rapidly selected for resistant lineages. Whole-genome sequencing revealed that four of five independently evolved clones carried mutations within the O9 O-antigen biosynthesis locus located adjacent to the CPS gene cluster (Fig. 5A, Fig. S2D). Two independent missense mutations were identified in *manC*, while a third clone carried a single-base deletion in *manB* that introduced a premature stop codon. (Table S3) ManB and ManC catalyze sequential steps in GDP-mannose synthesis, generating the activated mannose donor required for assembly of the O9 polymannan O-antigen (Fig. 5B) (48, 49). A fourth resistant isolate carried a single-base deletion in *wzt*, resulting in a premature stop codon. Wzt is a component of the ABC transporter responsible for translocating mature O9 polymannan chains across the inner membrane prior to ligation to the lipid A-core by WaaL (Fig. 5B) (48).

**Figure 5.**
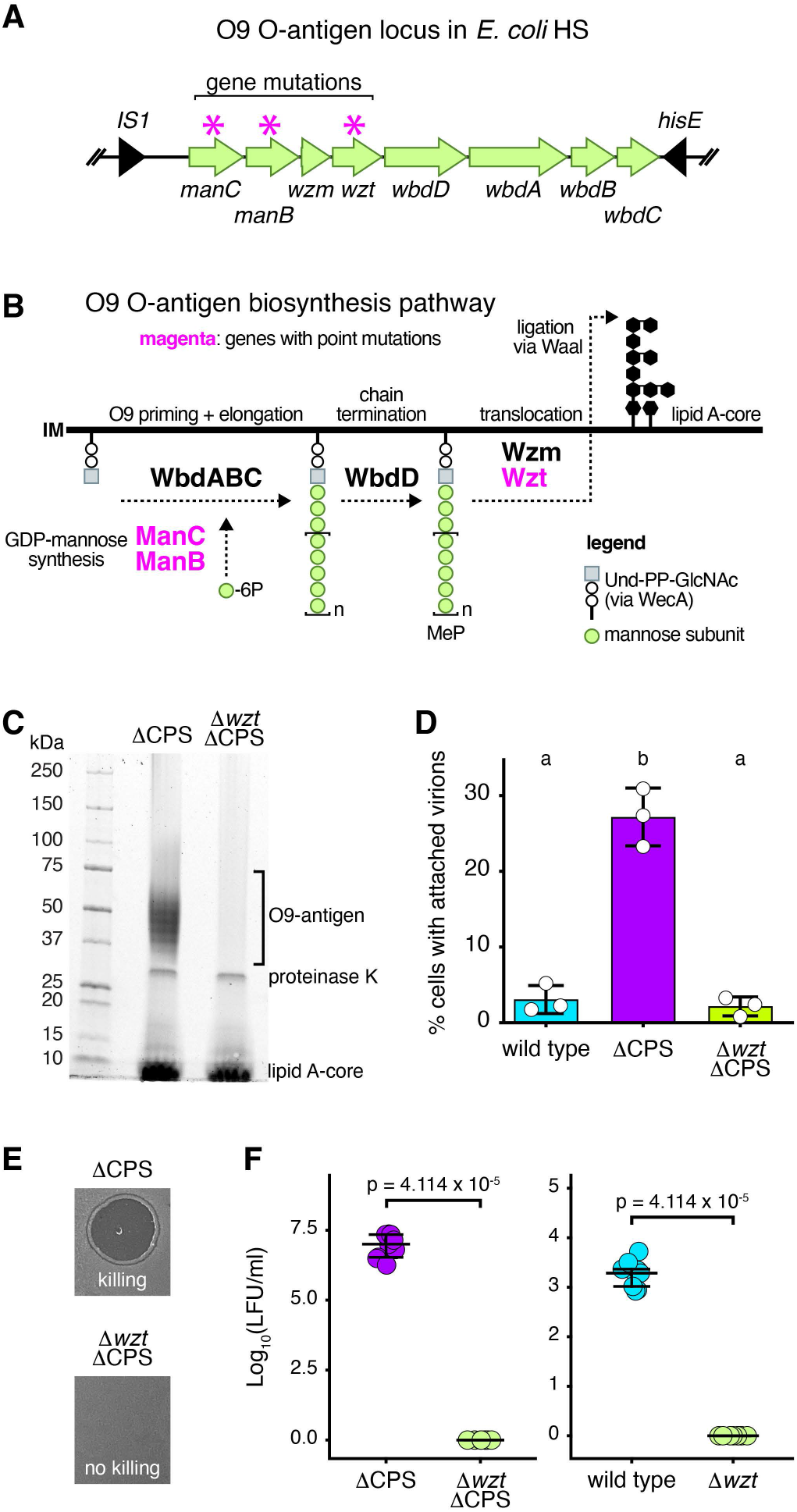
O9 O-antigen is required for DuoHS infection and cell killing. **(A)** Organization of the O9 O-antigen biosynthesis locus in *E. coli* HS. Green arrows denote genes involved in O9 O-antigen production. Magenta asterisks indicate genes carrying mutations in independently evolved DuoHS-resistant clones. **(B)** Predicted pathway for O9 O-antigen biosynthesis in *E. coli* HS. ManB and ManC generate GDP-mannose from mannose-6-phosphate, which provides the activated sugar donor for O9 polymannan synthesis by WbdA, WbdB, and WbdC on a WecA-dependent Und-PP-GlcNAc primer. Mature O-antigen chains are terminated by WbdD-mediated methyl phosphate (MeP) capping, exported across the inner membrane (IM) to the periplasmic space by the Wzm-Wzt ABC transporter, and ligated to the lipid A-core by WaaL. The resulting LPS is subsequently transported to the cell surface by the LPS transport (Lpt) system (not shown). Genes mutated in independently evolved DuoHS- resistant clones are highlighted in magenta. **(C)** Pro-Q Emerald-stained SDS-PAGE analysis of lipopolysaccharide isolated from ΔCPS and Δ*wzt* ΔCPS strains. The characteristic ladder-like O9 O-antigen banding pattern is seen in the ΔCPS strain and absent in the Δ*wzt* ΔCPS mutant, confirming loss of O9 O-antigen. **(D)** Percentage of cells carrying at least one attached DuoHS virion 15 min post-exposure to fluorescent Phollow phage virions. Wild-type and ΔCPS datasets are replotted from Fig. 3B for comparison. Loss of O9 O-antigen production (Δ*wzt*) in the ΔCPS background restores attachment frequencies to wild-type levels. Bars indicate means ± s.d. from three biological replicates (shown as individual points). Data were analyzed by one-way ANOVA followed by Tukey’s multiple-comparison test. Different letters indicate significant differences (p < 0.05). **(E)** Spot assay of an undiluted DuoHS lysate on ΔCPS and Δ*wzt* ΔCPS bacterial target lawns. Zones of clearing indicate phage-mediated killing, whereas confluent bacterial growth indicates resistance to phage attachment and/or infection. **(F)** Lysogen formation following DuoHS infection of ΔCPS and Δ*wzt* ΔCPS target strains (left) and wild-type and Δ*wzt* target strains (right), quantified as lysogen formation units (LFU/ml). Bars indicate the median and interquartile range across biological replicates (n = 9). Statistical significance was determined using Fisher’s exact test (p = 4.11 x 10^-5^).

To directly test the role of O9 O-antigen in DuoHS infection, we made targeted deletions of *wzt* in both wild-type and ΔCPS DuoHS target strain backgrounds. Pro-Q Emerald staining confirmed the absence of O9 O-antigen in the Δ*wzt* ΔCPS mutant (Fig. 5C). Unlike the parental ΔCPS strain, Δ*wzt* ΔCPS double mutants were highly resistant to DuoHS binding and killing (Fig. 5D,E). Loss of O9 O-antigen also abolished lysogenic infection of both ΔCPS and wild-type hosts (Fig. 5F). These findings identify O9 O-antigen as the host receptor required for DuoHS infection and suggest that capsule expression limits infection by occluding access to this underlying structure.

### Hyperion and Kronos states determine prophage–bacterium competitiveness and influence community assembly

We next examined how the balance between Hyperion and Kronos states within microbial communities influences coexistence between DuoHS prophages and their hosts. Here, we define coexistence as the mutual persistence of prophage and host populations following lytic outbreaks, when conflict between phage replication and host fitness is most pronounced. Based on our observation that capsule loss restricts phage dissemination while increasing host mortality, we predicted that Hyperion states would promote competitive coexistence by supporting prophage transmission without excessive host damage. In contrast, we predicted that Kronos states would compromise coexistence by increasing lethal outcomes while limiting prophage spread.

To test these predictions, we assembled defined two-member communities comprising equal mixtures of a lysogen population carrying a DuoHS prophage and a DuoHS-cured target population. Lysogen and target populations were independently established in either a Hyperion (wild-type, capsule-expressing) or Kronos (ΔCPS) state, generating all four possible combinations of host–phage interactions. Communities were briefly induced with MMC to trigger DuoHS lytic replication, then transferred to solid media prior to virion release, allowing spatial community assembly to occur during an active phage outbreak. Community composition and prophage transmission were quantified by fluorescence imaging, using mKate2-labeled target cells and a DuoHS-encoded mNeonGreen lysogeny reporter to identify transmitting lysogens and track prophage spread into surrounding target populations (Fig. 6A). This experimental setup enabled direct visualization of how Hyperion and Kronos states shape community structure during phage outbreaks (Fig. 6A,B).

**Figure 6.**
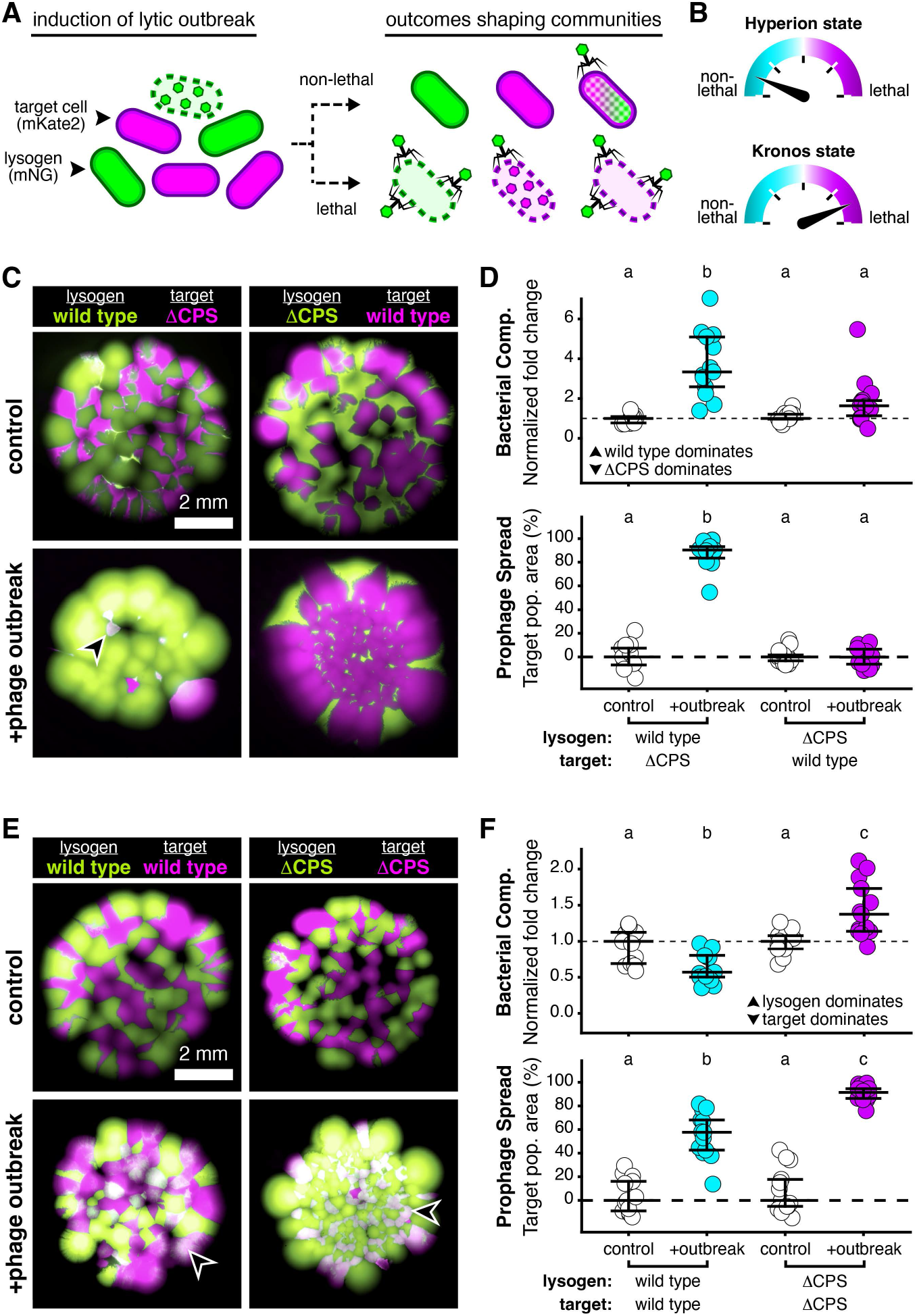
Hyperion and Kronos regimes control patterns of prophage–host coexistence in spatially structured communities. **(A)** Experimental scheme used to quantify the effects of prophage outbreaks on community assembly. Two-member communities consisted of DuoHS lysogens carrying an mNeonGreen (mNG) lysogeny reporter and mKate2-labeled target cells. Following induction of a lytic outbreak, non-lethal outcomes include lysogenic infection, generating double-labeled cells, or no detectable infection. Lethal outcomes include productive lytic replication within target cells and collateral killing of lysogens or target cells. **(B)** Conceptual model illustrating the predicted relative frequency of lethal and non-lethal interactions in Hyperion and Kronos states. Hyperion states favor non-lethal interactions, whereas Kronos states favor lethal interactions. **(C)** Representative images of interstrain communities composed of wild-type (Hyperion) lysogens and ΔCPS (Kronos) target cells (left), or ΔCPS lysogens and wild-type target cells (right). Communities are shown before (top) or after (bottom) induction of a prophage outbreak. White areas (arrowheads) indicate newly lysogenized target populations carrying both fluorescent markers. **(D)** Quantification of interstrain community outcomes from panel C. Upper panel: bacterial community composition before and after prophage outbreak induction. The dashed line denotes equal abundance of the two competing populations. Values above the line indicate enrichment of wild-type populations, whereas values below the line indicate enrichment of ΔCPS populations. Lower panel: prophage spread quantified as the percentage of target population area occupied by newly lysogenized cells. The dashed line at zero indicates the absence of detectable prophage transmission. **(E)** Representative images of intrastrain communities composed entirely of (left) Hyperion state populations (wild-type lysogens and wild-type targets) or (right) Kronos state populations (ΔCPS lysogens and ΔCPS targets). Communities are shown before (top) or after (bottom) prophage outbreak induction. White areas (arrowheads) indicate newly lysogenized target populations. **(F)** Quantification of intrastrain community outcomes from panel E. Upper panel: bacterial community composition before and after prophage outbreak induction. The dashed line denotes equal abundance of lysogen and target populations. Values above the line indicate lysogen enrichment, whereas values below the line indicate target cell enrichment. Lower panel: prophage spread quantified as the percentage of target population area occupied by newly lysogenized cells. The dashed line at zero indicates the absence of detectable prophage transmission. Bars in plots indicate the median and interquartile range. Points represent individual communities (n = 13 communities from seven biological replicates). White symbols denote uninduced control communities, whereas cyan and purple symbols denote communities experiencing outbreaks initiated by Hyperion-state and Kronos-state lysogens, respectively. Statistical significance was assessed using two-way ANOVA with community type and outbreak status as fixed effects, followed by Tukey-adjusted pairwise comparisons. Different letters indicate statistically significant differences (p < 0.05).

In the absence of induction, all community configurations remained balanced, with lysogen and target populations persisting at similar relative abundance and exhibiting minimal prophage spread (Fig. 6C–F, ‘control’). Thus, in the absence of prophage-mediated conflict, Hyperion and Kronos states had little effect on community composition. Following lytic induction, however, community outcomes diverged dramatically according to the states of the interacting populations.

When wild-type lysogens competed with ΔCPS targets, wild-type populations dominated community composition following the lytic outbreak, with DuoHS prophages spreading extensively into the few remaining pockets of capsule-deficient target cells (Fig. 6C,D).

Emphasizing a causal role for prophage attack in determining this outcome, we found that both an inducible DuoHS prophage in wild-type lysogens and O9 O-antigen expression in the ΔCPS target population were required (Fig. S5). In the reciprocal configuration, when ΔCPS lysogens competed with wild-type targets, capsule-deficient lysogens were strongly depleted in several replicate communities and prophage transmission into the capsule-expressing target populations remained near baseline levels (Fig. 6C,D). These experiments reveal a pronounced asymmetry in outbreak outcomes, with Hyperion lysogens simultaneously maintaining host fitness and promoting prophage dissemination, whereas Kronos lysogens exhibit reduced competitiveness and limited prophage spread.

Distinct assembly outcomes also emerged in communities where both lysogens and target populations occupied the same Hyperion or Kronos state. In wild-type Hyperion communities, lysogens experienced a modest fitness disadvantage, consistent with the partial loss of lysogens through the induction of lytic replication, while prophage spread into target populations remained moderate (Fig. 6E,F). In contrast, ΔCPS Kronos communities exhibited substantially greater prophage transmission but also favored lysogens over target cells, indicating that localized outbreaks suppress target population growth (Fig. 6E,F).

Altogether, these experiments begin to reveal how Kronos and Hyperion interactions can shape patterns of microbial community assembly. Consistent with our prediction, Hyperion states promoted prophage transmission and bacterial competitive fitness, whereas Kronos states undermined the persistence of one or both partners. However, the distinct outcomes observed in communities composed entirely of either Kronos or Hyperion populations suggest that these states are not simply binary interaction regimes, but instead define a continuum that, depending on community context, generates diverse patterns of host fitness, prophage spread, and community structure.

## Discussion

Mutualistic symbioses are shaped by conflict as much as by cooperation (50–52). Tension arises because each partner can benefit at the other’s expense, in some cases leading to collapse of the association. Persistence therefore depends on mechanisms that limit exploitation, align fitness interests, and maintain net benefits across ecological and evolutionary contexts (50–52). Through this lens, we sought to clarify the sources of conflict in prophage– bacterium symbioses (4, 5, 52). By experimentally evolving a prophage–bacterium association toward exploitation and antagonism, we gained broader perspectives on both the scale at which conflict emerges and the mechanisms that mitigate it.

First, prophage–bacterial conflict is often viewed at the level of individual cells, yet we identified an additional layer operating at the population level, where lytic outbreaks can generate locally high densities of virions capable of inflicting collateral damage on susceptible target cells and sister lysogens. Second, transitions between mutualistic and antagonistic states are typically considered to be under the control of phage-encoded regulatory programs governing lysogenic and lytic replication. However, we found that control can instead reside with the host, as capsule expression limits phage adsorption and prevents self-destructive collateral attack. This host-mediated constraint on autoinfectivity, which we term the Hyperion effect, is mutually beneficial because it promotes both host survival and phage transmission (Figs. 3, 4, and 6).

Our work builds upon the recently described Kronos effect, in which phage-encoded superinfection exclusion systems act as “anti-Kronos” factors by blocking reinfection of viral progeny (43). Capsule shielding achieves a similar outcome through host-mediated control of phage adsorption, highlighting how distinct host- and phage-encoded traits can preserve infectious virions and reduce autoinfectivity. Therefore, within the Kronos–Hyperion continuum, both phages and bacterial hosts can actively regulate the balance between conflict and cooperation.

Although the evolutionary history of the DuoHS–HS association is unknown, our findings suggest a plausible route by which Hyperion-like states may arise. In an ancestral capsule-deficient host permissive to DuoHS reinfection, self-destructive Kronos interactions would be expected to impose strong selection for reduced prophage adsorption and autoinfectivity. One possible evolutionary response is the acquisition of capsule biosynthesis genes, likely through horizontal gene transfer from *Klebsiella* or related taxa (32, 53), enabling host control over phage attachment. Consistent with this idea, natural *E. coli* Lambda lysogens exhibit not only intracellular immunity but also resistance to prophage binding, which has been proposed to reduce both infection by virulent phages sharing the same receptor and autoinfection by prophage progeny (46). Our findings further suggest that collateral damage arising from lysis from without may constitute an additional source of selection favoring surface-level resistance, particularly in dense communities where lytic outbreaks generate locally high concentrations of virions. While the precise mechanism by which capsule expression in HS restricts DuoHS attachment remains unresolved, these converging pressures suggest that Hyperion-like solutions could emerge through diverse strategies that limit phage adsorption, including receptor modification, surface shielding, or phase variation. More broadly, many previously characterized resistance mechanisms may influence where associations reside along the Kronos–Hyperion continuum.

In our model, host capsule production enhances the spread of DuoHS. By limiting Kronos-mediated consumption of phage progeny, host investment in a Hyperion state preserves infectious virions and promotes their dispersal. Although this may appear altruistic, our experiments in two-member communities show that prophage–host pairs in a Hyperion state outcompete those in a Kronos state (Fig. 6). In this context, phages can be viewed as weapons deployed by their hosts to eliminate competitors (4, 52). Phage dispersal thus becomes a competitive strategy operating at the level of the phage–bacterium partnership. Our findings extend previous observations that prophage induction in a subset of cells can enhance the overall fitness of a lysogen population during competition (54). Importantly, this advantage depends on bacterial traits regulating phage infection. Here, receptor compatibility and capsule shielding together determine whether phage-mediated competition can occur, acting to define both the targets and beneficiaries of prophage transmission. In addition, our findings, together with recent work in chitin-based biofilms of *Vibrio cholerae*, highlight that prophage-mediated competition is a spatially regulated process, contingent on the local arrangement and density of lysogens, susceptible hosts, and dispersing virions (55).

An important question that emerges from our work is how Kronos–Hyperion interactions differ among prophage–bacterium symbioses and what factors govern their outcomes. Based on the P2-like phage DuoHS studied here and observations from natural *E. coli* Lambda lysogens, we expect analogous interactions may be common (46). However, their strength and consequences likely vary. For example, in the JBD26–*Pseudomonas* prophage–host association from which the Kronos effect was originally described, reinfection does not appear to harm lysogens, suggesting that the fitness consequences of Kronos interactions can vary considerably (43). One interpretation is that destructive reinfection could be averted by multiple layers of Hyperion-like protections that limit the fitness consequences of autoinfectivity. In line with this idea, our community assembly experiments suggest that prophage–bacterium partnerships may occupy different positions along a Kronos–Hyperion continuum rather than being locked into discrete states (Fig. 6 and Fig. S5). From this perspective, Kronos and Hyperion interactions do not simply determine whether associations persist or collapse. Instead, depending on the mechanisms involved, they may tune the balance of fitness tradeoffs experienced by both bacterial hosts and their prophage symbionts, generating diverse outcomes in transmission, competitiveness, and ultimately community structure.

Evidence from other systems suggests that Kronos–Hyperion dynamics may extend beyond temperate phages. A quantitative study of a marine bacterial system indicates that cellular debris generated during lytic phage replication can inhibit or sequester infectious virions (56) Under these conditions, the accumulation of debris was proposed to slow phage propagation and prevent host extinction, thereby stabilizing the association. This Kronos-like feedback parallels our finding that, in the DuoHS–HS system, lysis-derived material also acts to dampen viral infectiousness. However, whereas we find that Hyperion interactions appear to stabilize prophage–bacterium symbioses by limiting autoinfection, Kronos-like interactions may promote coexistence in lytic phage systems. Future comparative studies will be needed to determine whether the stability of associations depends more strongly on ecological context or phage life cycle.

Finally, it remains to be explored whether Kronos–Hyperion interactions between individual phage–host pairs might scale to influence complex communities. Although our findings are derived from a minimal model system comprising a single phage–host pair under defined in vitro conditions, they provide a starting point for considering this possibility. In our system, capsule expression functions as a Hyperion safeguard by limiting collateral prophage attack; however, recent work shows that predatory lytic phages can infect HS by targeting its K47 capsule (32), highlighting a costly tradeoff that could drive ongoing ecoevolutionary dynamics with consequences for bacterial strain diversity and virome composition. Similar questions emerge from systems such as *Bacteroides thetaiotaomicron*, where phase-variable capsules generate heterogeneous populations that differ in phage susceptibility and contribute to long-term coexistence between bacterial hosts and the virulent phages that prey upon them (56) In such systems, considering potential roles for Kronos–Hyperion interactions may reveal additional processes governing coexistence within complex microbiomes, including those of the vertebrate gut. More broadly, because bacterial capsules are known to affect both microbe-microbe and animal-microbe interactions, Kronos–Hyperion mechanisms modulating bacterial phenotypes may extend their influence beyond phage–bacterium interactions alone.

## Acknowledgements

We are grateful to Ami Bhatt (Stanford University) for her generosity with both time and expertise. Early discussions about mobile genetic elements, sequencing-based approaches for studying them, and the broader goals of this project were invaluable in shaping its direction and development.

We are grateful to the community of the Marine Biological Laboratory course *Molecular and Cell Biology of Symbiosis* for insightful discussions that influenced several of the broader questions considered in this work. We especially thank Rose Faucher (University of California, Irvine), Derek Wu (Emory University), Anna Golikova (University of Miami), Austin Link (University of Oklahoma), Joselyn Molinar (Indiana University), Raluca Hedes (University of Freiburg), and Phillip Cleves (University of California, Berkeley) for their generosity with time, energy, and thoughtful conversations.

This work was supported by the National Institute of Allergy and Infectious Diseases of the National Institutes of Health under Award Number DP2AI154420 (to T.J.W.); the U.S. Army Research Office under Grant Number W911NF2510185 (to T.J.W.); the National Institute of General Medical Sciences under Award Number R35GM142797 (to S.G.); the Office of the Director and the National Center for Complementary and Integrative Health of the National Institutes of Health under Award Number DP2AT013749 (to M.E.G.); the University of California Office of the President Research Grants Program Office under award R00RG2814 (to K.L.W.); and the National Science Foundation Graduate Research Fellowship Program under Grant Number DGE-2235784 (to J.T.L.). M.Z. was supported by a PostDoc.Mobility fellowship from the Swiss National Science Foundation (P500PB_214357). The content is solely the responsibility of the authors and does not necessarily represent the official views of the National Institutes of Health. Any opinions, findings, conclusions, or recommendations expressed in this material are those of the authors and do not necessarily reflect the views of the NIH, NSF, Army Research Office, or the U.S. Government. The U.S. Government is authorized to reproduce and distribute reprints for Government purposes notwithstanding any copyright notation herein.

This study was made possible in part through access to the Optical Biology Core Facility of the Developmental Biology Center, a shared resource supported by the Cancer Center Support Grant CA-62203, the National Institutes of Health grant NIH S10OD032327-01, and the Center for Complex Biological Systems Support Grant GM-076516 at the University of California, Irvine.

## Author Contributions

Conceptualization: M.Z., L.O.d.O., T.J.W.

Methodology: M.Z., L.O.d.O., T.J.W.

Investigation: M.Z., L.O.d.O., J.T.L., Y.J.C., Ma.E.G., K.H., T.J.W.

Formal Analysis: M.Z., L.O.d.O., T.J.W.

Visualization: M.Z., L.O.d.O., T.J.W.

Resources: N.H.-R., E.T.W., T.P.O.

Supervision: T.J.W., S.G., M.E.G., K.L.W.

Funding Acquisition: T.J.W., S.G., M.E.G., J.T.L., M.Z.

Writing – Original Draft: M.Z., T.J.W.

Writing – Review & Editing: M.Z., T.J.W., L.O.d.O., M.E.G., K.L.W.

## Competing interests

The authors declare no competing interests.

**Movie S1.** DuoHS transmission and secondary lytic replication in a ΔCPS target cell.

**Movie S2.** Collateral prophage attack leading to lysis from without in a ΔCPS target cell

**Table S1.** Low-frequency SNP mutations in the CPS locus and signals of structural variations in the CPS locus.

**Table S2.** IS1 elements in *E. coli* HS

**Table S3.** Mutations recovered in O9 O-antigen biosynthesis genes

**Table S4.** Bacterial strains, primers, and plasmids

## Methods

### Bacterial Strains, Phages, and Culture Conditions

All wild-type and recombinant bacterial and phage strains used or generated in this study are listed in Table S4. Bacterial stocks were maintained at −80°C in 25% glycerol. For routine use, bacterial strains were inoculated directly from frozen stocks into 5 ml Tryptic Soy Broth (TSB; MP Biomedicals, VWR) and grown overnight (∼16 h) at 37°C with shaking. For bacterial cultivation on solid media, tryptic soy agar (TSA; Hardy Diagnostics, VWR) was used. Where indicated, growth media were supplemented with gentamicin (10 μg ml^-1^), chloramphenicol (20 μg ml^-1^), or anhydrotetracycline (aTc; 50 ng ml^-1^). The temperate bacteriophage DuoHS and derivative prophage variants were used throughout this study. DuoHS is a naturally occurring, DNA damage-inducible P2-like prophage integrated within the chromosome of *Escherichia coli* HS (18, 31, 58). Lytic induction of DuoHS was routinely performed using MMC treatment where indicated. Prophage variants carrying fluorescent reporters, antibiotic resistance markers, or SpyTag modifications were constructed and used as described below.

### Phollow Terminology

Fluorescence-based tracking of DuoHS induction, dispersal, and transmission was performed using the previously described Phollow system (18). Throughout this study, *E. coli* HS strains carrying SpyTag-modified DuoHS prophages and expressing fluorescent SpyCatcher proteins are referred to as “Phollow lysogens”. These strains were previously described as Phollow virocells (18); however, because the present study focuses on prophage–bacterium interactions, we use the term lysogen throughout. During lytic replication, fluorescent SpyCatcher proteins accumulate on and covalently bind assembling DuoHS capsids, generating correspondingly labeled “Phollow phages” or “Phollow phage virions”. Phollow lysogens are identified according to the fluorescent SpyCatcher protein they express (e.g., mNeonGreen or mKate2 Phollow lysogens), whereas Phollow phage virions are identified according to the fluorescent label acquired during replication in the host cell from which they originated. Unless otherwise indicated, “target cells” refer to DuoHS prophage-cured recipient populations that are susceptible to DuoHS infection and permissive to either lytic or lysogenic replication.

### Quantification of DuoHS Replication, Transmission, and Infection

#### DuoHS prophage induction and virion purification

DuoHS induction was done by treating *E. coli* HS lysogens with a pulse of MMC (Goldbio) as previously described (18). Briefly, overnight cultures of *E. coli* HS lysogens were diluted 1:100 into fresh TSB and grown for 1 h at 37°C with shaking. MMC was then added to the culture at a final concentration of 8 μg ml^-1^ for 1 h. Following this pulse treatment, cells were pelleted by centrifugation at 10,000 x *g* for 2 min and washed twice with 0.7% (w/v) NaCl (Sigma-Aldrich), resuspended in fresh 5 ml TSB, and incubated for an additional 2 h at 37°C with shaking to allow DuoHS lytic replication and release. To harvest virions, cultures were centrifuged at 21,000 x *g* for 1 min, and the supernatant with released virions was collected and treated with 0.1x volume of chloroform (Sigma-Aldrich) for 10 min. The mixture was centrifuged at 21,000 x *g* for 3 min to achieve phase separation. The aqueous phase containing phage virions was recovered and stored at 4°C for up to 3 days before use.

#### Quantification of released DuoHS virions by qPCR

The number of assembled and intact DuoHS virions (i.e., encapsidated viral DNA) was quantified as a proxy for released virions by quantitative PCR (qPCR). This was accomplished by extracting DuoHS DNA using the Monarch Genomic DNA Purification Kit (New England Biolabs) following the protocol for Gram-positive bacteria, with minor modifications. To deplete free DNA, lysates were treated with DNase I (0.2 U; BIOSEARCH Technologies) for 60 min at 37°C, followed by enzyme inactivation with 5 mM EDTA at 75°C for 10 min prior to DNA extraction. DNA was eluted in DNase- and RNase-free water using a volume equivalent to the original lysate volume. To specifically quantify the number of DuoHS genomes, primers MZP06 and MZP07 flanking the DuoHS attachment site (*attP*) were used. qPCR reactions were performed with PowerUp SYBR Green Master Mix (Applied Biosystems) on a CFX Duet Real-Time PCR System (Bio-Rad) using the following cycling conditions: 50°C for 2 min, 95°C for 2 min, and 40 cycles of 95°C for 15 s and 60°C for 1 min. Absolute viral DNA copy numbers were determined from a standard curve generated from serial dilutions of purified phage DNA. Cycle threshold (Ct) values were plotted against the known DNA copy number and fitted by linear regression, and the resulting equation was used to calculate phage DNA copy numbers.

#### Quantification of released DuoHS virions by lysogen-forming units

The number of DuoHS particles capable of lysogenic infection was quantified by lysogen-forming units (LFU) as previously described (18). Briefly, an overnight culture of a gentamicin-resistant *E. coli* HS ΔDuoHS target strain was diluted 1:400 into fresh TSB and grown for 30 min at 37°C with shaking, then washed twice and suspended in 0.7% (w/v) NaCl. The optical density at 600 nm was measured, and cultures were adjusted to 10^8^ colony-forming units (CFU) ml^-1^ in 0.7% (w/v) NaCl. As the infecting phage, an engineered DuoHS variant conferring chloramphenicol resistance was used. Phages were added to the target strain suspension at a multiplicity of infection (MOI) of 10, and cultures were supplemented with 5 mM CaCl^2^ (Ward’s Science) and incubated for 1 hour at 37°C with shaking. To enumerate lysogens, infected cultures were plated on TSA supplemented with chloramphenicol and gentamicin and incubated for 18 hours before colonies representing newly formed lysogens were counted.

#### Growth response during antibiotic stress

To assess the growth responses of the wild-type and ΔCPS strains to MMC and polymyxin B sulfate (PMB; GoldBio), overnight cultures were grown in 5 mL TSB at 37°C with shaking. Overnight cultures were diluted 1:10^6^ into fresh TSB and cultured for an additional 3 h under the same conditions. Cultures were then diluted 1:3 into TSB containing MMC at 0–16 μg ml^-1^ or PMB at 0–32 μg ml^-1^. Aliquots were dispensed into sterile, clear-bottom 96-well microplates, and optical density at 600 nm was recorded over time using a BMG Labtech FLUOstar Omega plate reader controlled by Omega software v5.7.

Concentrations exceeding the minimal inhibitory concentration were excluded from the visual representation of growth curves.

### Genetic Engineering of Bacterial and Phage Strains

#### Tn7-mediated chromosomal insertions

Insertion of fluorescent markers or SpyCatcher variants, antibiotic resistance genes, or genetic switches into the *attTn7* site of the bacterial chromosome was performed using a pTn7xTS transposon-based system (59). The tagging vector pTn7xTS and transposase-encoding vector pTNS2 (Addgene #64968) were introduced into *E. coli* HS by either electroporation or triparental mating using *E. coli* SM10 donor strains. For mating reactions, donor and recipient strains were mixed at a 1:1:1 ratio, spotted onto a filter disk placed on TSA, and incubated at 30°C for ∼4 h. Cells were then recovered and plated onto TSA supplemented with gentamicin, followed by overnight incubation at 37°C to select for insertion variants. Tn7 insertion at the *attTn7* site adjacent to *glmS* gene (MJ392_21845) was confirmed by PCR using primers WP11 and WP150.

#### Construction of capsule (ΔCPS) mutants

CPS-deficient mutants lacking the four-gene locus targeted by the IS1A element (*wzb* (MJ392_08255)*, wzc* (MJ392_08260),*wbaP* (MJ392_08265), and a glycosyltransferase-encoding gene ( MJ392_08270)) were generated using two independent approaches: λ Red-mediated recombination (60) or markerless allelic exchange (59). For the λ Red-derived mutants, the kanamycin resistance gene from plasmid pKD4 was amplified using primers MZP57 and MZP58, each containing 40-bp overhangs homologous to the 5’ and 3’ end of the deletion region. The resulting amplicon was introduced by electroporation into *E. coli* HS strains carrying pKM208 vector, encoding an isopropyl-β-d-thiogalactopyranoside (IPTG)–inducible λ Red recombinase. Recombinants in which the kanamycin cassette replaced the four-gene locus were selected on TSA supplemented with kanamycin (50 μg ml^-1^) and verified by PCR using primers MZP59 and MZP52. For markerless deletion, the four-gene CPS locus was deleted using the pAX1 allelic exchange as previously described (59). An allelic exchange cassette was constructed by amplifying the 5’ homology region (925 bp, primers MZP60 and MZP61) just upstream of *wbaP* and the 3’ homology region (850 bp, primers MZP62 and MZP63), which included the glycotransferase stop codon and downstream region. The two homology arms were inserted into the *Sma*l-linearized pAX1 vector using in vivo assembly (IVA), and correct assembly was confirmed by PCR (primer WP106 and WP204) and whole-plasmid sequencing (Plasmidsaurus). The resulting vector, pMZ252, was used for markerless deletion of *wbaP*, *wzb*, *wzc*, and glycosyltransferase genes by allelic exchange. Deletion of the locus was confirmed using primers MZP59 and MZP52.

#### Construction of O9 O-antigen (Δ*wzt*) mutants

Markerless deletion of the *wzt* gene (MJ392_08350) within the O9 O-antigen locus was performed using the pAX1 allelic exchange system as previously described (59). An allelic exchange cassette was constructed by amplifying a 5’ homology region (974 bp, MZP66 and MZP67) including the start codon of *wzt* and the upstream region and the 3’ homology region (862 bp, primers MZP64 and MZP65), which included the *wzt* stop codon and downstream region. The two homology regions were inserted into the *Sma*l-linearized pAX1 vector using IVA, and correct assembly was confirmed by PCR (primer WP106 and WP204) and whole-plasmid sequencing (Plasmidsaurus). The resulting vector pMZ253 was used for markerless deletion of *wzt* by allelic exchange. Correct deletion of the locus was confirmed using primers MZP68 and MZP69.

#### Construction of marked DuoHS prophage variants

##### Chloramphenicol-resistant DuoHS

To quantify lysogenic infectivity via cultivation methods, a chloramphenicol resistance cassette was inserted into the second moron region of the DuoHS prophage, located between the tail genes *gpH* (MJ392_14345) and *gpFI* (MJ392_14340). The resistance gene was amplified from plasmid pKD3 using primers containing 40 bp homology arms flanking the target integration site. WP356 and WP357 were used when inserting in the wild-type DuoHS prophage and primers MZP42 and WP357 were used when inserting into DuoHS prophages carrying the mNeonGreen lysogeny reporter (described below). PCR amplicons were then introduced into *E. coli* HS carrying the IPTG-inducible λ Red recombination plasmid pKM208. Recombinants were selected on chloramphenicol-containing media and verified by PCR using primers WP358 and WP357.

##### mNeonGreen DuoHS lysogeny reporter

To visually follow DuoHS lysogenic infection, a gene encoding mNeonGreen was inserted into the second moron region of DuoHS prophage by allelic exchange. A custom pUC57 plasmid (GenScript) containing homology arms corresponding to the DuoHS tail fiber and major tail sheath genes was used as an intermediate cloning vector. The mNeonGreen expression cassette, driven by the synthetic P^tac^ promoter, was excised from pTTW49 by EcoRI and AvrII digestion, blunted, and ligated between the DuoHS homology arms. Correct insertion was confirmed by PCR and Sanger sequencing using primers MZP36 and MZP37. The mNeonGreen cassette and flanking DuoHS homology arms were subsequently excised from the intermediate plasmid by restriction digestion and ligated into the pAX1 allelic exchange vector, generating pAX1-DuoHS-moron::mNG (pMZ84). Correct assembly was confirmed by PCR using primers MZP38 and MZP39. The vector pMZ84 was used to introduce the P^tac^-driven mNeonGreen cassette into the second moron region of DuoHS by allelic exchange, and correct integration (DuoHS moron::mNG) was confirmed by PCR using primers WP358 and WP359.

#### Design and construction of inducible capsule and lysis switches

##### Capsule switch construction

The previously constructed gain-of-function switch (pTn7xTS-GOF-switch, pTW285) was used as the backbone (61). To generate the ΔCPS^iCPS^ strain, the contiguous genes *wzb* (MJ392_08255), *wzc* (MJ392_08260), *wbaP* (MJ392_08265), and a glycosyltransferase gene (MJ392_08270) were amplified using primers MZP53 and MZP54, including 5’ and 3’ homology overhangs to the delivery vector. The PCR amplicon was inserted into the cargo site of the pTn7xTS-GOF-switch vector using IVA, creating pMZ201. Correct insertion was confirmed by complete plasmid sequencing (Plasmidsaurus). The verified pMZ201 vector was used to insert the inducible gain-of-function switch into markerless **Δ**CPS strains.

##### Lysis switch construction

The DNA sequence encoding the phage λ lysis genes *S*, *R*, and *Rz* was obtained from the λ genome (GenBank accession no. J02459.1). A synthetic DNA fragment containing the *SRRz* locus, including the native ribosome binding sites associated with each gene, was synthesized by GenScript (Piscataway, NJ) and cloned into a pUC57 backbone flanked by SnaBI restriction sites, generating pTW220. The *SRRz* cassette was excised from pTW220 by SnaBI digestion and ligated into the SmaI cargo site of pTn7xTS-GOF-switch (61), producing pTW297. pTW297 was introduced into *E. coli* SM10 by electroporation, generating strain TW303. The lysis switch was delivered into wild-type and ΔCPS *E. coli* HS strains by Tn7- mediated transposition using established methods (59). Lysis switch function was validated using disk diffusion test in which overnight cultures of *E. coli HS* carrying the lysis switch was spread onto a TSA. Sterile filter disks were placed onto bacterial lawns and loaded with either sterile water or aTc. Plates were incubated overnight at 37°C and subsequently imaged using a Leica MZ10 F fluorescence stereomicroscope equipped with 1.0x and 1.6x objectives, GFP and Texas Red filter sets, and a Leica K5 camera (Leica, Wetzlar, Germany). Images were processed using standard Leica Application Suite software and FIJI (62). For visualizing the extracellular release of dTomato protein, 1 ml from overnight cultures with and without aTc- induction was centrifuged at 21,000 x g for 1 min to pellet cells in a 1.6 ml tube. Tubes were illuminated with a green (520–535 nm wavelength) LED flashlight and imaged with an iPhone 13 Pro affixed with Roscolux (#342 Rose Pink) filter sheet.

### Experimental Evolution of the DuoHS–HS Symbiosis

#### Serial induction and reinfection

DuoHS prophages carrying a chloramphenicol resistance marker were induced as described above, with the modification that, in the first passaging experiment, lysogens were continuously exposed to MMC for 16 hours prior to lysate harvest. A 50 μl aliquot of lysate was added to a 1:100-diluted overnight culture of an *E. coli* HS ΔDuoHS target strain (5 ml total volume) and incubated for 8 h at 37°C with shaking. Aliquots of 100 μl were then plated on tryptic TSA supplemented with gentamicin and chloramphenicol to select for de novo lysogens. All colonies obtained on each plate were harvested and stored as cryostocks, which served as the inoculum for the subsequent passage. The second passaging scheme followed the same workflow, except that phage induction was performed as described in “DuoHS prophage induction and virion purification,” and virion abundance in the lysate was quantified to normalize the MOI to 10. Infections were carried out in 0.7% (w/v) NaCl supplemented with 5 mM CaCl^2^ for 1 hour at 37°C with shaking, followed by plating on TSA supplemented with gentamicin and chloramphenicol. In the second passaging scheme, 20 colonies per plate were collected to initiate the next cycle.

### Genomic Analysis of Evolved Lysogens

#### DNA extraction, sequencing, and quality control

Bacterial DNA from de novo lysogens was extracted using the Wizard Genomic DNA Purification Kit (Promega) following the manufacturer’s protocol for Gram-negative bacteria. DNA quality and quantity were assessed using a NanoDrop spectrophotometer (260/280 = 1.89; 260/230 = 2.06) and a DeNovix DS-11 Fx fluorometer. For library preparation, 40 ng of DNA was processed using the Nextera DNA Flex Library Prep Kit (Illumina) following the low-volume protocol (63). Sequencing was performed at the University of California, Irvine Genomics Research and Technology Hub (GRT-Hub, Irvine, CA, USA) on a NextSeq 200 platform, generating 2 × 150 bp paired-end reads with a targeted insert size of 300 bp and an output of approximately 8 million reads per sample. Raw sequencing data were quality-controlled using *bbduk.sh* from BBMap (v.38.71). Adapter sequences were removed, followed by filtering of reads mapping to the PhiX genome. Low-quality reads were discarded using the parameters *trimq=14*, *maq=20*, *maxns=1*, and *minlength=45*.

#### Variant identification by Breseq

Breseq (v0.38.3) was used to detect genetic and structural variants relative to a curated reference genome (64, 65). The reference genome used was *E. coli* HS (NCBI accession CP092639). To minimize strain-specific variation, the reference was polished using quality-controlled, trimmed reads from the wild-type *E. coli* lysogen. Detected variants were incorporated into the reference genome using *gdtools APPLY* with GenBank-formatted output using the flag *-f GENBANK* for subsequent read mapping and variant calling. For the initial passaging experiment, *breseq* was run in consensus mode (*-c*), and in the second passaging experiment, in polymorph mode (*-p*) to enable detection of minority variants within a community. For each sample, the output was inspected in the breseq HTML report.

#### Detection of structural variants by MGEfinder

To detect genome rearrangements arising from mobile genetic element (MGE) insertions, we utilized MGEfinder (v1.0.6) following the developer’s recommended workflow (24). Raw Illumina paired-end reads were first deduplicated using *hts_SuperDeduper* (HTSStream toolkit), then subjected to quality and adapter trimming with *trim_galore* (parameters: --fastqc --paired). Trimmed reads were aligned to the reference genome (CP092639) using BWA-MEM (v0.7.8), and alignment files were converted and formatted for downstream analysis using the built-in MGEfinder *formatbam* command (66). Additionally, quality-controlled reads were used as input for de novo assembly of lysogen genomes using SPAdes (v.3.15.4, parameter *-isolate*) (67). At last, the MGEfinder was executed in de novo workflow mode (*mgefinder --workflow –denovo*) to identify mobilizable elements.

#### Verification of IS1A insertion by read alignment

To verify IS1A insertion into the *wbaP* gene of clone 2 (passage 1), the clone was subjected to Oxford Nanopore sequencing, and de novo assembly done by SeqCoast Genomics, LLC (Portsmouth, NH, USA). Samples were lysed in MagMAX Microbiome Bead Beating Tubes with Qiagen CD1 lysis buffer using a Vortex-Genie 2 for 5 min. The library was prepared using the Oxford Nanopore Native Barcoding Kit (SQK- NBD114), sequenced on a PromethION 2 Solo, and base-called using the Dorado super-accurate model. Quality-controlled reads were used for de novo assembly with Unicycler (68). Illumina short reads were aligned to the de novo assembly using BWA-MEM (v.0.78).

Unmapped, secondary, and supplementary alignments were excluded, and primary alignments with a mapping quality ≥20 were retained, sorted, and indexed using SAMtools (69). Reads mapping to the IS1A target region of contig 1 (1,732,967–1,744,600 bp) were extracted and converted to FASTQ format using bamToFastq (BEDTools). The extracted reads were realigned to the corresponding locus reference using BWA-MEM. The resulting alignments were sorted, and the base-pair coverage was calculated using SAMtools depth (-a) and then visualized in R (70).

#### PCR-based detection of IS1A insertion

Primers MZP59 and MZP52 flanking the capsule biosynthesis locus were used to detect IS1A insertions by PCR. Reactions were performed using Q5 polymerase (Invitrogen), and amplicons were resolved on 0.8% (w/v) agarose gels at 7.5 V cm^-1^ for 5 h. IS1A insertion events were identified by the expected 768 bp shift in amplicon size relative to the wild-type locus.

### Capsule and O-Antigen Characterization

#### Capsule and O-antigen biosynthesis gene prediction and comparative genomics

The capsule type and biosynthesis genes of *E. coli* HS strain were predicted using the webpage Kaptive (71), designed to predict and classify the capsule in *Klebsiella* species. The O-antigen type and biosynthesis genes were predicted using SerotypeFinder v2 with an ID threshold of 0.85 and a minimum length of 0.6 (72). The amino acid sequences of the following reference genomes (CP092639.1, CP102490.1, CP122617.1, CP064678, and CP007799.1) were downloaded from NCBI for homology comparisons and visualization with pyGenomViz using MUMmer settings (73).

#### Capsule Staining

Wild-type, HS*^IS1A^*^::*wbaP*^, ΔCPS, and ΔCPS^iCPS^ strains were streaked on TSA or grown overnight in TSB at 37°C. For ΔCPS^iCPS^ strains, cultures and plates were additionally supplemented with aTc, (Clontech) to induce capsule production. Exopolysaccharide production by *E. coli* HS was visualized using tannin–mordant staining as previously described (74). Colonies were smeared onto microscopy slides, heat-fixed, and stained with fuchsin solution (0.3 g basic fuchsin in 10 mL 95% ethanol, brought to 100 ml with 5% phenol) for 3 min. Slides were rinsed, heat-dried, and treated with a freshly prepared tannin–mordant solution (2 volumes 0.3 g L^-1^ FeCl^3^, 2 volumes 1.5 g L^-1^ tannins, 5 volumes 2 g L^-1^ saturated potassium aluminum sulfate) for 3 min, then washed. For counterstaining, smears were exposed to 1% methylene blue for 30 s. Samples were imaged with an Echo Revolve microscope using a 60x oil-immersion objective; exopolysaccharides appeared blue and cells red.

#### Cell pellet compaction assay

To assess cell compaction phenotypes, 1 ml of overnight culture was pelleted at 7000 × *g* for 2 min, washed three times with Milli-Q water, and imaged with a digital camera.

#### LPS extraction and Pro-Q staining

Approximately 1 billion cells of *E. coli* ΔCPS or ΔCPS Δ*wzt* were pelleted at 4000 × *g*, washed with 1 ml phosphate-buffered saline (PBS), and suspended in a final volume of 1 ml PBS. Aliquots of 50 μl were pelleted and suspended in 2x Laemmli Buffer (Bio-Rad). Cells were lysed by boiling for 10 min, followed by cooling to room temperature. Proteinase K (8000 units ml^-1^, New England Biolabs) was added to the lysate and incubated overnight at 55°C. Proteinase K was subsequently inactivated by boiling for 10 min. Aliquots of 10 μl were separated on a 4-15% Criterion TGX Protein Gel (Bio-Rad) at 200 V for 40 minutes. The gel was stained with Pro-Q Emerald 300 (Invitrogen) according to the manufacturer’s instructions and imaged using the Pro-Q channel on a ChemiDoc MP Imaging system (Bio-Rad).

#### Selection for DuoHS-resistant ΔCPS variants

To identify the putative receptor mediating DuoHS attachment and killing, ΔCPS lysogens were challenged with high titers of purified DuoHS virions to select for resistant variants. Overnight cultures of wild-type (control) and ΔCPS lysogens were diluted 1:10^6^ in TSB and grown for 3 h at 37°C with shaking. The culture was then diluted 1:3, mixed 1:1:2 with TSB and DuoHS lysate, and incubated for 24 h at 37°C with shaking. Bacterial growth was monitored by measuring the optical density at 600 nm. Cultures with visible growth were subjected to a second round of DuoHS exposure. Cells were diluted 1:100 in fresh TSB, grown for 3 h, then diluted 1:200 into a TSB-phage lysate mixture containing two parts lysates. Cultures were incubated for an additional 24 hours under the same conditions. After each round of phage exposure, aliquots were spread onto TSA plates using sterile cotton swabs, and 10 μl of phage lysate was spotted onto each bacterial lawn to assess resistance. Following two rounds of enrichment, six independently derived resistant ΔCPS lysogen lineages were selected for whole-genome sequencing at SeqCoast Genomics, LLC (Portsmouth, NH, USA) on a NextSeq200 platform, generating 2 × 150 bp paired-end reads. Data processing and breseq analysis to identify mutations (**Table S3**) were performed as described above.

### Imaging Phage Induction and Transmission with Phollow

#### Tracking adsorption and lytic replication with purified DuoHS Phollow phage virions

To measure virions adsorption, we used purified mNeonGreen Phollow phage virions and mKate2- SpyCatcher-expressing wild-type, ΔCPS, or Δ*wzt*ΔCPS target strains. Target strains were grown overnight in TSB at 37°C with shaking, diluted 1:25 into 5 ml fresh TSB, and incubated for 30 min at 37°C with shaking. Following the incubation, the entire culture was pelleted by centrifugation at 21,000 for 1 min and suspended in 1 ml of 0.7% (w/v) NaCl. 150 μl of the culture was then transferred into five individual 1.6 ml tubes and supplemented with 5 mM CaCl^2^. 15 μl of lysate containing Phollow phage virions was added to each tube and incubated at 37°C with shaking for 0, 15, 30, 45, or 60 min. At each time point, a single tube was centrifuged at 21,000 × g for 1 min to collect cells and washed twice with 20 μl 0.7% (w/v) NaCl before being resuspended to a final volume of 20 μl. The phage/bacteria suspension was then combined 1:1 with 1% low-melt agarose (Sigma), and 1 μl of the mixture was mounted on a glass slide and sealed with a coverslip. Fluorescence imaging was performed on an Echo Revolve microscope equipped with a 60x oil-immersion objective and GFP and Texas Red filter sets. Multiple non-overlapping fields of view (3–9) were acquired for each biological replicate. For each field of view, the total number of mKate2-SpyCatcher expressing bacterial cells, the number of cells with one or multiple bound mNeonGreen Phollow phage virions and the number of cells undergoing secondary lytic replication were manually counted. Bound virions were classified as those that were overlapping or directly adjacent to a bacterial cell. Phage attachment frequency (%) was calculated as (number of cells with attached virions/total cells counted) x 100. Each biological replicate (n = 3) consisted of an independent overnight culture and an independent phage lysate sample.

#### Imaging DuoHS reinfection during induced phage outbreaks

Overnight cultures of wild- type and ΔCPS mNeonGreen Phollow lysogens were diluted 1:100 into 5 ml fresh TSB and grown for 1 h at 37°C with shaking. Cultures were treated for 1 h with a pulse of MMC (8 μg ml^-1^), before being pelleted by centrifugation at 10,000 x *g* for 2 min, washed twice with 0.7% (w/v) NaCl (Sigma-Aldrich), and suspended in a final volume of 5 ml TSB. Cultures were placed back at 37°C with shaking and imaged 1 h post-wash by fluorescence microscopy on an Echo Revolve microscope equipped with GFP and Texas Red filter sets and a 60x oil-immersion lens.

#### Time-lapse imaging of DuoHS transmission

To visualize DuoHS prophage induction and transmission by time-lapse microscopy, cultures of wild-type mNeonGreen Phollow lysogens and ΔCPS target cells expressing mKate2-SpyCatcher were first grown separately overnight in 5 ml TSB at 37°C with shaking. The following day, cultures were diluted 1:100 into fresh TSB and grown for an additional hour under identical conditions. Equal volumes (60 μl) of each culture were then combined and transferred to an 8-well #1.5 high-precision slide chamber (Cellvis). Prior to inoculation, wells were coated with 0.1% poly-D-lysine (Thermo Scientific) to promote cell immobilization. Chambers were centrifuged at 1,400 × *g* for 15 min at room temperature to facilitate bacterial attachment to the bottom of the well, after which the supernatant was removed and wells were washed three times with 200 μl of 0.7% saline to remove non-adherent cells. Lytic replication was induced by addition of 150 μl TSB supplemented with 0.2% low-melt agarose (Sigma) and 8 μg ml^-1^ MMC. Chambers were incubated for 5 min at room temperature to allow agar solidification before imaging on a Zeiss Elyra 7 microscope. Time-lapse images were acquired at 20-min intervals using structured illumination microscopy, and 488 nm and 560 nm laser lines to detect mNeonGreen and mkate2, respectively.

#### Image Processing and Analysis

Raw images were processed in Fiji (62) as previously described for Phollow imaging (18). Adjustments to brightness and contrast were applied across images intended for visualization. For Echo Revolve images capturing virion adsorption and lytic replication, the built-in Fiji *Despeckle* and *Smooth* functions were applied to improve image clarity for presentation in figures. Custom lookup tables (LUTs) used to visualize cells and virions were identical to those described previously for Phollow imaging (18). Unless otherwise indicated, image processing was not used to alter or influence quantitative measurements.

### Virion Entrapment and Dissemination Assays

#### Cell-mediated virion pull-down

To measure levels of virion adsorption, purified wild type-derived DuoHS virions carrying a chloramphenicol resistance gene were added to 1 ml of 0.7% (w/v) NaCl containing 10^8^ CFU ml^-1^ of either the wild-type or ΔCPS target strain. As a negative control, phages were added to 1 ml of 0.7% NaCl without bacterial cells. All mixtures were incubated at 37°C with shaking for 1 h to allow virion adsorption, then centrifuged at 21,000 x *g* for 2 minutes. The resulting supernatant was then used to quantify unbound, free phage by mixing it with gentamicin-resistant wild-type target cells (10^8^ CFU ml^-1^). This mixture was incubated at 37°C with shaking for 1 h. After incubation, cells were plated on TSA supplemented with gentamicin and chloramphenicol to select for and quantify LFUs.

#### Inhibition of infectivity with cell-free lysis debris

To determine whether host material released during lysis could reduce phage infectivity, cell-free lysates were generated from wild- type or ΔCPS strains engineered to express λ lysis genes under the control of an aTc-inducible promoter. Overnight cultures were diluted 1:100 into TSB supplemented with 50 ng ml^-1^ aTc and incubated for 24 h to induce lysis. Cultures were then passed through a 0.45-μm syringe filter (EMD Millipore) to remove intact cells and collect cell-free lysates. Gentamicin-resistant ΔCPS target cells were resuspended in the indicated cell-free lysates and infected with purified chloramphenicol-resistant DuoHS virions at an MOI of 100. Following infection, cultures were plated on TSA supplemented with gentamicin and chloramphenicol to quantify LFU. In this assay, a ΔCPS target strain was used for LFU quantification because its heightened susceptibility to DuoHS infection provides greater sensitivity for detecting small differences in lysogenic infectivity.

#### Virion dissemination assay

To quantify the spatial dissemination of infectious phage particles through bacterial communities, mKate2-SpyCatcher-labeled wild-type and ΔCPS target populations were mixed at ΔCPS:wild-type ratios of 100:0, 50:50, 10:90, and 1:99. Cell mixtures were washed twice in 0.7% (w/v) NaCl and diluted 1:100 into 0.7% (w/v) tryptic soy soft agar prior to overlaying onto TSA plates. Prior to applying the agar overlay, a sterile 1 ml pipette tip was placed onto the TSA plate (tip side up) to cast a central well within the soft-agar overlay. DuoHS virions carrying the mNeonGreen lysogeny reporter were mixed 1:1 with molten 1.4% (w/v) tryptic soy soft agar and deposited into the central well. Plates were incubated at 37°C for 24 h and imaged using a Leica MZ10 F fluorescence stereomicroscope equipped with 1.0x and 1.6x objectives, GFP and Texas Red filter sets, and a Leica K5 camera (Leica, Wetzlar, Germany). Images were processed using standard Leica Application Suite software and FIJI (62). For the standardized virion dissemination assay, 60 mm plates containing a 5 ml TSA base layer and a 3 ml overlay of 0.7% soft agar were used. Soft agar was prepared by adding UltraPure Agarose (Invitrogen) to TSB at a final concentration of 1.4 or 7% (w/v) before sterilization.

As infectious virions disseminate and propagate through the surrounding bacterial population, newly lysogenized target cells acquired green fluorescence, generating a visible dissemination front. Dissemination was quantified in Fiji (ImageJ), and the distance from the central inoculation site to the leading edge of the dissemination front was measured at three independent positions for each replicate community, and the mean value was used as the dissemination distance for that replicate.

### Competitive Fitness and Survival Assays

#### Head-to-head lysogenic infection

Relative virion fitness was quantified using pairwise lysogenic infection competitions between DuoHS particles derived from different host backgrounds. DuoHS virions derived from capsule mutants (ΔCPS or HS^IS1::*wbaP*^) were mixed 1:1 with wild-type-derived virions, yielding a final concentration of 10^9^ phage ml^-1^. As a control, wild-type-derived virions were used as the competitor. Virion abundance was normalized by qPCR prior to mixing. One phage competitor in each pair carried a fluorescent lysogeny reporter, allowing the origin of newly established lysogens to be determined by colony fluorescence. Following infection, LFUs were enumerated, and the ratio of competitor-derived to wild-type-derived LFUs was Log^10^-transformed to compare relative lysogenic fitness.

#### Spot Assays

Wild-type and ΔCPS, with and without DuoHS prophages, were grown overnight, washed twice in 0.7% (w/v) NaCl, and diluted 1:100 in 0.7% (w/v) tryptic soy soft agar before overlaying onto TSA plates. Purified wild-type-derived DuoHS virions carrying an mNeonGreen lysogeny reporter were serially diluted and 3 μl of each dilution was spotted onto the dried soft-agar overlays. Plates were incubated at 37°C and imaged using a Leica MZ10 F fluorescence stereomicroscope equipped with 1.0x and 1.6x objectives, GFP and Texas Red filter sets, and a Leica K5 camera (Leica, Wetzlar, Germany). Images were processed using standard Leica Application Suite software and FIJI (62). Soft agar was prepared by adding UltraPure Agarose (Invitrogen) to TSB to a final concentration of 0.7% (w/v).

#### DisCo-like spot assays

Fluorescently labeled mKate2 wild-type and mNeonGreen ΔCPS lysogen strains were grown overnight, mixed at a 1:1 ratio, and washed twice in 0.7% (w/v) NaCl. The cell mixture was diluted 1:100 in 0.7% (w/v) tryptic soy soft agar and overlaid onto TSA plates. 3 μl of purified wild-type-derived DuoHS virions were spotted onto the dried soft-agar overlays, and plates were incubated at 37°C overnight prior to being imaged using a Leica MZ10 F fluorescence stereomicroscope equipped with 1.0x and 1.6x objectives, GFP and Texas Red filter sets, and a Leica K5 camera (Leica, Wetzlar, Germany). Images were processed using standard Leica Application Suite software and FIJI (62). Soft agar was prepared by supplementing tryptic soy broth with 0.7% (w/v) of UltraPure Agarose.

#### Lysogen competitions

Fluorescently labeled ΔCPS, HS^IS1A::*wbaP*^, and wild-type lysogens, and an unlabeled wild-type strain, were grown overnight in TSB at 37°C with shaking. Each fluorescent lysogen was mixed 1:1 with the unlabeled wild-type strain, washed twice with 0.7% (w/v) NaCl, and diluted 1:100 into TSB. Cultures were pulse treated with MMC and incubated for 24 h. Samples were taken at indicated time points to enumerate abundances of each population. Uninduced controls were handled identically, except that MMC was omitted, and samples were collected at the beginning and after 24 h. The ratio of competitor to wild-type CFUs for each pairwise competition was Log^10^-transformed and plotted to compare relative abundances.

#### Two-member community Assembly Experiments

Overnight cultures of mKate2-labelled wild-type or ΔCPS target strains were mixed 1:1 with either a wild-type or ΔCPS lysogen, carrying a DuoHS prophage with an mNeonGreen lysogeny reporter. Mixtures were washed twice with 0.7% (w/v) NaCl and suspended in an equal volume of TSB. Each mixture was then used to inoculate a 5 ml TSB culture at a 1:100 dilution. To confirm equal starting abundances, serial dilutions were prepared in 0.7% (w/v) NaCl, and a total volume of 10 μl per dilution was applied as six separate spots of roughly 1.6 μl each onto agar plates. Plates were incubated at 37°C for 24 h, and strains were quantified by counting CFU based on their fluorescent markers; images of 1:100 dilution spots were acquired as pre-induction “controls”. The remaining culture was then pulse-induced with MMC (6 μg ml^-1^). After antibiotic washout, undiluted and 1:10 diluted cultures were spot-plated onto TSA. Plates were incubated at 37°C for 24 h, and imaged using a Leica MZ10 F fluorescence stereomicroscope equipped with 1.0x and 1.6x objectives, GFP and Texas Red filter sets, and a Leica K5 camera (Leica, Wetzlar, Germany). Images were processed using standard Leica Application Suite software and FIJI (62).

Community composition and prophage dispersal were quantified from fluorescence images that were thresholded to generate binary masks using Fiji (62). A custom macro using built-in image-calculation and area-measurement functions quantified the areas occupied by the different strains. Prophage spread was quantified as the spatial overlap between the two masks using a pixel-wise AND operation. The lysogen-specific area was calculated by subtracting the overlapping area from the total lysogen area. The target strain area, lysogen area, and overlapping area were measured separately and exported as a CSV file for further analysis and plotting in R.

#### Negative-stain Electron Microscopy (EM)

Samples were negative stained based on a previously published method (75). Gilder 200-mesh copper grids (Ted Pella) were floated on collodion film (Electron Microscopy Sciences), dried and carbon coated using a Leica ACE200. Prior to staining, all grids were negatively glow-discharged using a PELCO easiGlow. 3 μl of each sample was applied to the grid surface and adsorbed for ∼15 seconds. Excess sample was blotted using filter paper (Whatman). All grids were washed twice with 50 μl drops of Milli-Q water, blotted after each wash, followed by staining using two 50 μl drops of 0.75% Uranyl Formate. Each final drop of excess stain was blotted using filter paper and the grid was air dried. A JEOL-JEM 2100F 200kV transmission electron microscope, equipped with a Gatan OneView 4k x 4k camera, was used for imaging the samples. The resulting micrographs were saved in .dm4 format to retain image metadata for processing in Fiji. The addition of scale-bars and contrast-enhancement to the micrographs was done using Fiji (62).

#### Numerical Data visualization and Statistical Analysis

All experiments were performed with at least three biological replicates. Statistical analyses and data visualization were performed in R (v4.5.2) using RStudio (2025.9.2.41) with the *tidyverse* (v 2.0.0), *rcompanion* (v2.5.2) and *multcompView* (v0.1-11) packages(76–79). Depending on whether the assumptions of parametric testing were met, statistical significance was assessed using either one-way analysis of variance (ANOVA) followed by Tukey’s honestly significant difference (HSD) post hoc test or Kruskal–Wallis tests followed by pairwise Mann–Whitney U tests with Benjamini–Hochberg false-discovery-rate correction. Statistical groups are indicated by different let ters (p < 0.05).

---

**Figure S1.**
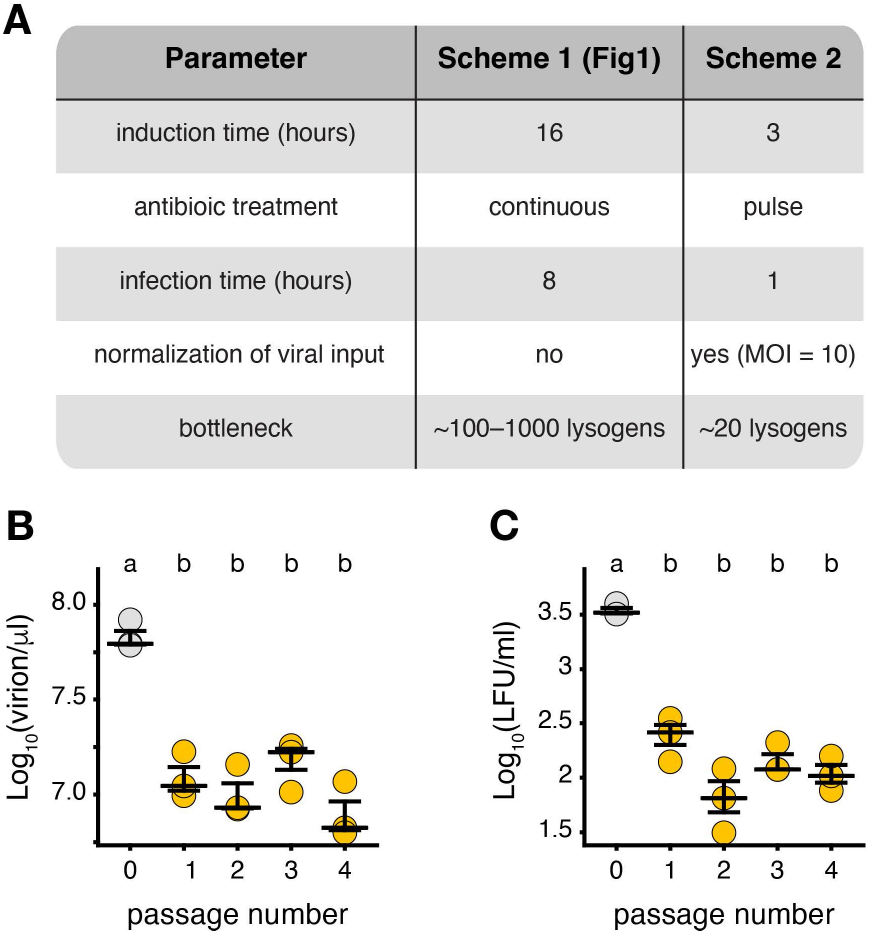
Experimental passaging parameters do not drive the decline in free DuoHS virions and lysogeny capacity. **(A)** Comparison of parameters used in the original (Scheme 1) and modified (Scheme 2) passaging experiment. For Scheme 2, the induction and infection durations were shortened, and fewer cells were used as an inoculum, while antibiotic exposure was limited to a pulse to induce lytic replication, and viral input was standardized to a multiplicity of infection (MOI) of 10. **(B)** Free virion yields measured by qPCR and **(C)** infectivity measured as lysogen-forming units (LFU) across serial passages. Grey circles represent data derived from ancestral lysogen (passage 0), whereas amber circles represent data derived from the lysogens archived at the indicated passages. Bars in each plot indicate the median and interquartile range. Statistical groupings denoted by different letters (p < 0.05) were determined by one-way ANOVA followed by Tukey’s HSD post hoc test for multiple comparisons.

**Figure S2.**
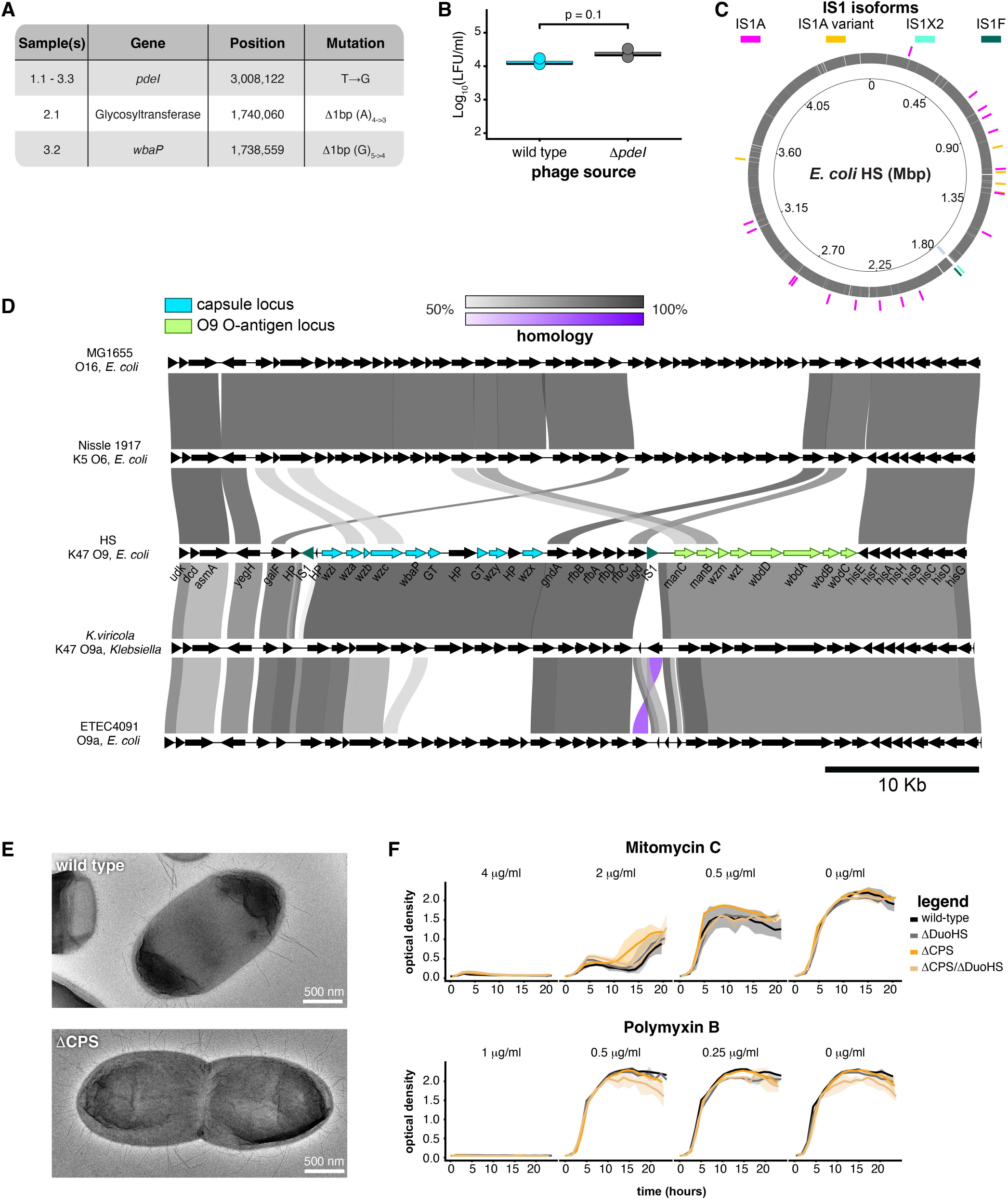
Genomic architecture and phenotypic characterization of capsule-deficient *E. coli* HS. **(A)** Mutations identified in de novo lysogens recovered during serial passaging. The table indicates the affected isolates, targeted gene, genomic position, and corresponding mutation. **(B)** Lysogenic infectivity of DuoHS in wild-type and Δ*pdeI* recipient strains, quantified as lysogen formation units (LFU). Bars indicate the median and interquartile range (n = 3). Statistical significance was assessed using a Mann–Whitney U (Wilcoxon rank-sum) test. **(C)** Circular genome map of *E. coli* HS. The chromosomal capsular polysaccharide (CPS) biosynthesis locus is highlighted in cyan in the inner ring, and genomic positions of IS1 isoforms are indicated on the outer ring. **(D)** Protein-level homology of the CPS and O9 O-antigen biosynthesis loci in *E. coli* HS relative to representative laboratory K12 (*E. coli* MG1655), commensal/probiotic (*E. coli* Nissle 1917), pathogenic (*E. coli* ETEC4091), and *Klebsiella* strains. CPS and O9 O-antigen biosynthesis genes in *E. coli* HS are highlighted in cyan and green, respectively. Shaded connectors indicate homologous proteins, with shading intensity corresponding to amino acid sequence identity. **(E)** Transmission electron micrographs of wild-type and ΔCPS *E. coli* HS cells. No obvious differences in cell morphology or extracellular polysaccharide structures were observed between the two strains. Scale bars, 500 nm. **(F)** Growth of wild-type and capsule-deficient strains in the presence of mitomycin C (top) or polymyxin B (bottom) measured by optical density (600 nm). Wild-type lysogen, wild-type ΔDuoHS, ΔCPS lysogen, and ΔDuoHS ΔCPS target strains displayed similar susceptibility across the tested concentrations. Lines indicate the mean, and shaded regions indicate the standard deviation (n = 3).

**Figure S3.**
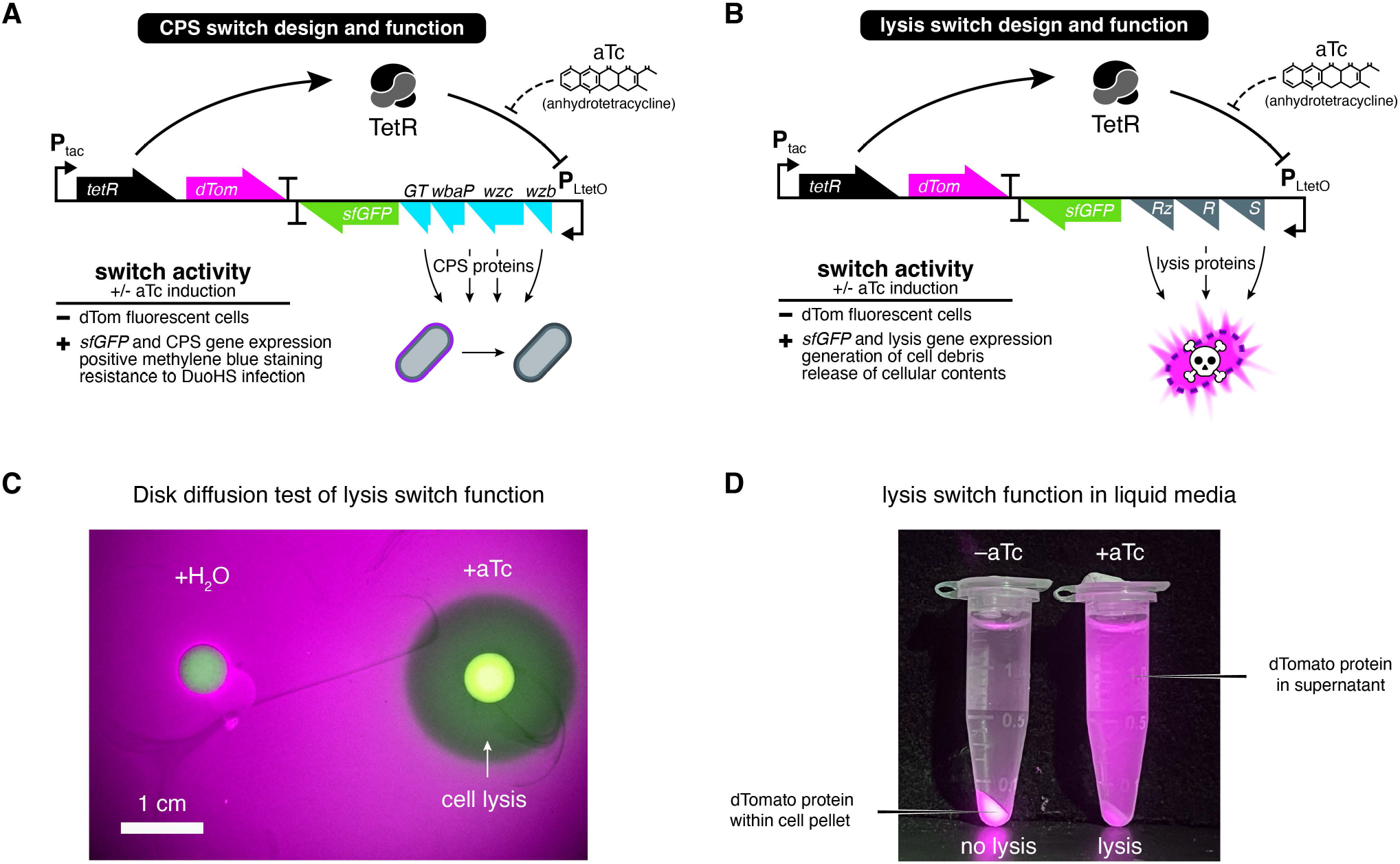
Design and functional validation of inducible capsule and lysis switches. **(A)** Schematic of the aTc-inducible capsule (CPS) switch. In the absence of anhydrotetracycline (aTc), constitutively expressed TetR represses the P^LtetO^ promoter, while constitutive dTomato (dTom) expression labels all bacterial cells. Addition of aTc relieves TetR-mediated repression, inducing expression of *sfGFP* and *wzb*, *wzc*, *wbaP*, and CPS glycosyltransferase (GT) genes. Capsule production results in positive methylene blue staining and resistance to DuoHS infection. sfGFP serves as a reporter of switch activation. **(B)** Schematic of the aTc-inducible lysis switch. As in panel A, aTc induction relieves TetR-mediated repression of P^LtetO^, resulting in expression of *sfGFP* and the λ phage lysis genes (*R*, *Rz*, and *S*). Induction causes cell lysis and release of cellular contents. **(C)** Disk diffusion assay demonstrating aTc-dependent activation of the lysis switch. A zone of clearance forms around a disk loaded with aTc (+aTc), whereas lysis is absent around the water control (+H^2^O) **(D)** Validation of lysis switch function in liquid culture using the release of cytoplasmic dTomato fluorescent protein as a readout. In the absence of aTc (-aTc), dTomato remains within intact cells and is retained in the cell pellet. Following aTc-induction (+aTc), dTomato is released into the supernatant as a consequence of lysis.

**Figure S4.**
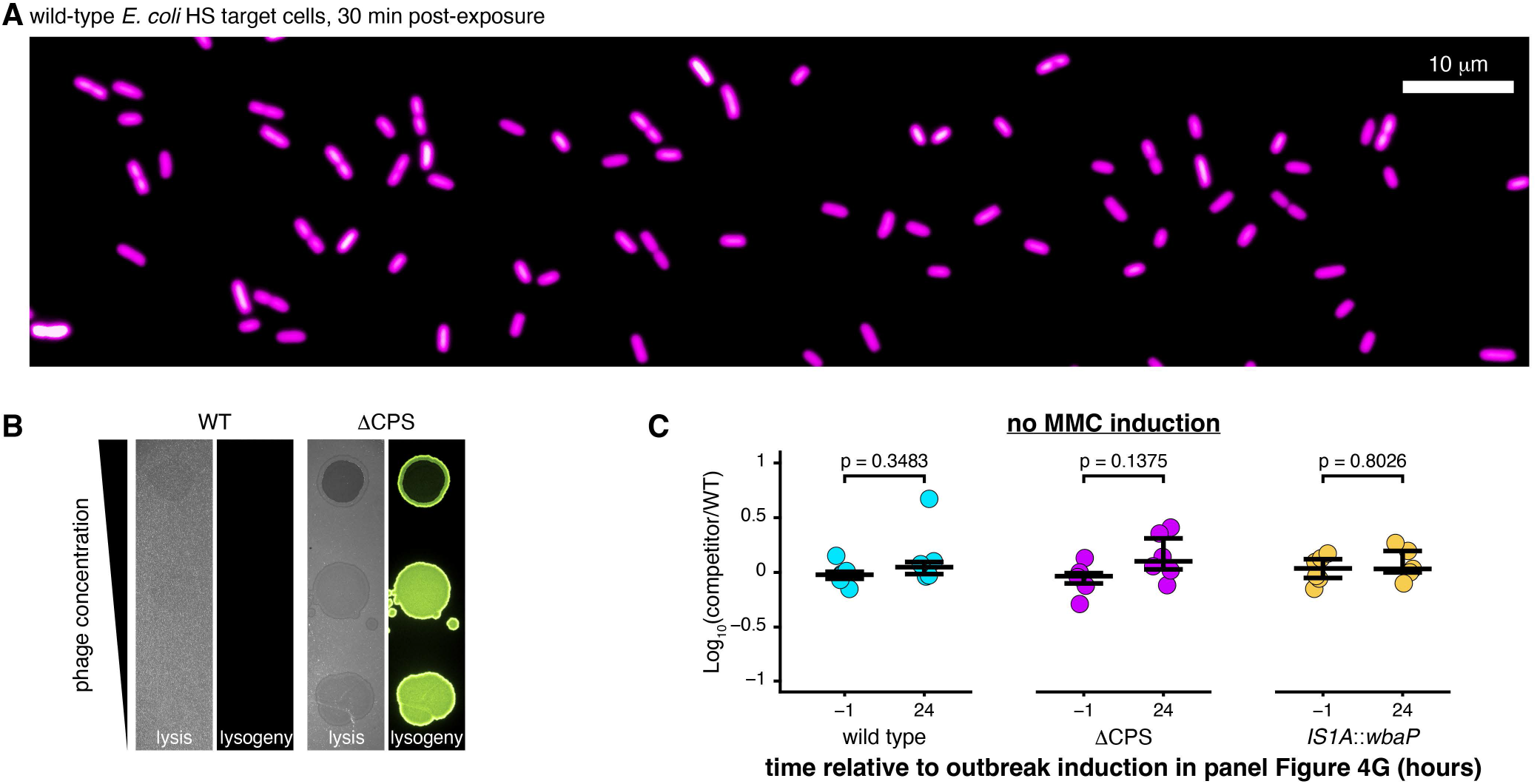
Control experiments supporting capsule-dependent protection against DuoHS attack. **(A)** Representative image of a wild-type *E. coli* HS target population 30 min following exposure to purified DuoHS Phollow phage virions. The image was acquired using the same DuoHS virion preparation, time point, and assay conditions shown in Fig. 4Diii. Unlike ΔCPS populations, widespread reductions in cytosolic fluorescence were not observed. Only the mKate2 fluorescence channel is shown. **(B)** Representative spot assays comparing wild-type and ΔCPS target populations exposed to wild- type-derived DuoHS virions carrying an mNeonGreen lysogeny reporter. Phage spots correspond to undiluted lysate and 1:32 and 1:500 dilutions (top to bottom). Brightfield images (left) show cell clearing, whereas fluorescence images (right) show mNeonGreen lysogeny reporter activity. Assays were performed as in Fig. 4E. **(C)** Relative abundance of capsule-deficient lysogens in the absence of MMC-induced prophage outbreaks. Wild-type lysogens were competed against either wild-type (cyan, n = 6), ΔCPS (purple, n = 6), or HS*^IS1A^*^::*wbaP*^ (amber, n = 6) lysogens. Relative abundance is expressed as Log^10^(CFU competitor/wild type). No significant differences were detected over 24 h in the absence of MMC treatment, indicating that the fitness differences observed in Fig. 4G are not attributable to intrinsic growth differences. Statistical comparisons were performed using two-sided paired t-tests.

**Figure S5.**
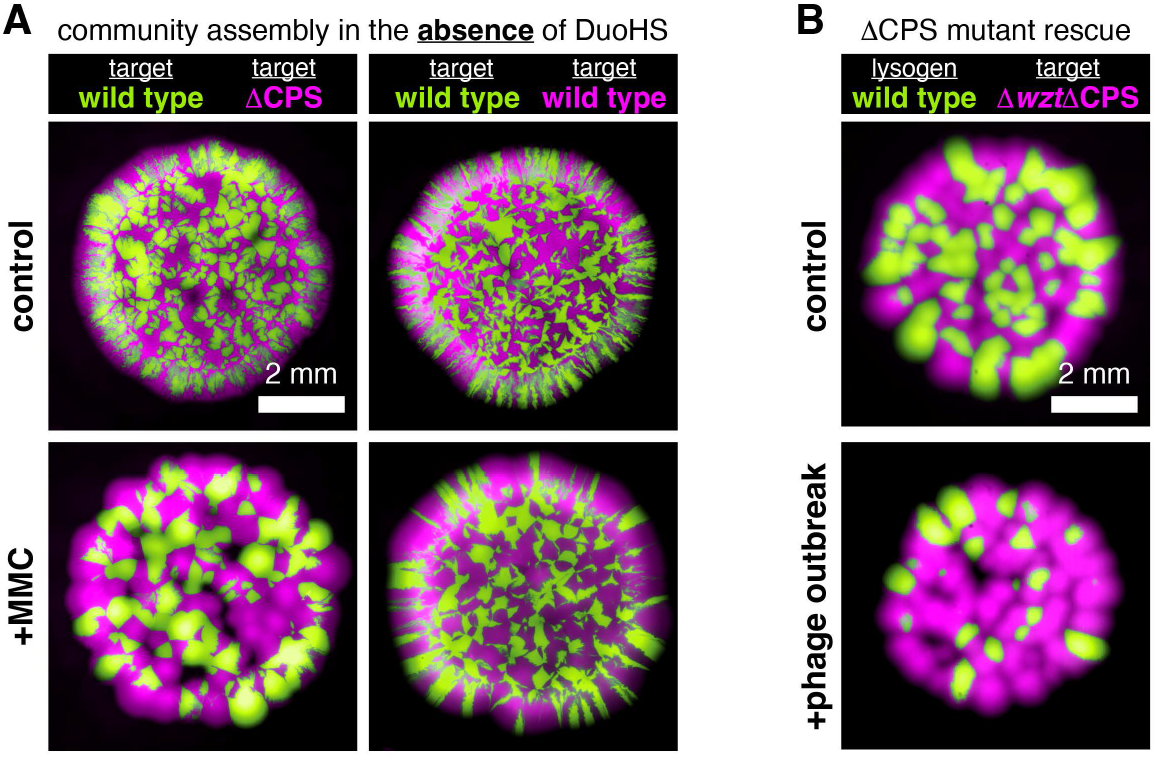
Community restructuring requires both DuoHS prophage carriage and O9 O- antigen expression. **(A)** Representative images of community assembly outcomes in the absence of a DuoHS prophage. Left: mixed target cell community composed of wild-type (Hyperion) and ΔCPS (Kronos) populations. Right: community composed entirely of wild-type (Hyperion) target populations. Communities are shown before (top) or 24 h after (bottom) MMC exposure. In the absence of a DuoHS prophage, MMC treatment alone does not reproduce the community restructuring observed during prophage outbreaks in Fig 6C/E. **(B)** Representative images of interstrain communities comprising a wild-type (Hyperion) lysogen and a ΔwztΔCPS target population. Communities are shown before (top) or after (bottom) induction of a prophage outbreak. Loss of the O9 O-antigen receptor in the target population prevents the extensive prophage spread and target cell depletion observed when wild-type lysogens compete with O9-expressing ΔCPS targets.

